# Identifying effective evolutionary strategies for uncovering reaction kinetic parameters under the effect of measurement noises

**DOI:** 10.1101/2024.03.05.583637

**Authors:** Hock Chuan Yeo, Vijay Varsheni, Kumar Selvarajoo

## Abstract

The transition from explanative modelling of fitted data to the predictive modelling of unseen data for systems biology endeavors necessitates the effective recovery of reaction parameters. Yet, the relative efficacy of optimization algorithms in doing so remains under-studied, as to the specific reaction kinetics and the effect of measurement noises. To this end, we simulate the reactions of an artificial pathway using 4 kinetic formulations: generalized mass action (GMA), Michaelis-Menten, linear-logarithmic, and convenience kinetics. We then compare the effectiveness of 5 evolutionary algorithms (CMAES, DE, SRES, ISRES, G3PCX) for objective function optimization in kinetic parameter hyperspace to determine the corresponding estimated parameters. We quickly dropped the DE algorithm due to its poor performance. Baring measurement noise, we find CMAES algorithm to only require a fraction of the computational cost incurred by other EAs for both GMA and linear-logarithmic kinetics yet performing as well by other criteria. However, with increasing noise, SRES and ISRES perform more reliably for GMA kinetics, but at considerably higher computational cost. Conversely, G3PCX is among the most efficacious for estimating Michaelis-Menten parameters regardless of noise, while achieving numerous folds saving in computational cost. Cost aside, we find SRES to be versatilely applicable across GMA, Michaelis-Menten, and linear-logarithmic kinetics, with good resilience to noise. On the other hand, we could not identify the parameters of convenience kinetics using any algorithm. Together, we find algorithms that are effective under marked measurement noise for specific reaction kinetics, as a step towards predictive modelling for systems biology endeavors.

## Introduction

In the book *A Brief History of Intelligence* [1], the author, Max S. Bennett, makes the important point of having a model that explicitly reproduces the fundamental features and interactions of a phenomenon (i.e., first-principled), to effectively learn the best course of action regarding it. For instance, the newly acquired perception of time and physical space by vertebrae has endowed a more advanced form of reinforcement learning, based on a model of the physical world. The new capability enables the schooling of sophisticated maneuvers, related to hunting and predator avoidance, as well as navigational skills, to enhance survival fitness in a harsh prehistoric world. Early mammals further acquire the capacity to flexibly simulate and hence discern alternative actions in their world model, providing them with a critical advantage over other animal classes, as to the efficiency and safety of learning. To note, the ancient competitive advantage of using a first-principle model for learning remains relevant in the current era of the fourth industrial revolution.

In an earlier commentary, we have set out the critical role of explicitly modelling the network and reaction kinetics for effective learning about biological systems [2]. For metabolic engineering [3], synthetic biology [4], or even precision drug endeavors [5], the holistic modelling of physico-chemical interactions among molecular components (systems biology [6]) is necessary for the reconstruction and elucidation of emergent behaviors and the identification of system-wide mechanisms and effects. Furthermore, the mathematical model of the change in component levels with time based on such interactions (i.e., rate laws or reaction kinetics), can be used to flexibly simulate, and predict component dynamics under broad conditions. From an application perspective, the mechanistic model also provides a structure grounded in first principles to help reduce sample size and datapoint requirement for training, effect of biological and technical noises, data leakage, as well as batch effects [2]. The model also grounds predictions in molecular mechanism, thereby allowing for experimental verification, and enabling explainability. Furthermore, in recent years, there has been increasing interest in achieving the aspiration of digital twin [7] in various endeavors [3–5]. Regardless, however, its underlying principle compels the mirroring of a physical twin in its dynamic behavior, which is only truly compatible with an explicit model describing the underlying pertinent mechanisms. In all, the mechanistic model remains central to any machine learning/artificial intelligence approach for effective learning about biological networks.

While systems biology has begun its nascent transition from explanatory modelling of fitted data to the predictive modelling of unseen data [2, 8], there remains two broad objectives in the field:

1. The learning of a mechanistic model of the system for predicting unobserved dynamics, a shift from the earlier, more restricted objective of calibrating the parameters of a given model to best reproduce the fitted data [9];
2. The finding of an optimal set of system input variables, given the design objective(s) for the biological system. This is typically done by repeatedly simulating the system, according to values sampled from the input variable hyperspace. The design landscape is then separately modelled and searched to find the set of input variables corresponding to the optimal design objective.

In this regard, researchers are cognizant of the challenges in learning such a mechanistic model, both in determining the underlying rate laws and their parameter values, which are sensitive to biochemical contexts, such as pH, temperature, ion concentrations, and the intracellular microenvironment [10–15]. A suggested approach to bypass the difficulty is to model the individual reaction rates as a black box ML model [8] but would result in a lack of clarity as to their biochemical basis. A more common heuristic approach is to first work on the selection or discernment of rate laws, followed by estimation of the associated parameters. Here, the point to note is that there remains room for improvement regarding the latter task in attaining the aspiration of predictive modelling for systems biology endeavours. The former task of determining the underlying form of the rate laws, and how to better integrate both tasks, presents further challenges that are not within the scope of the current study.

Currently, the various techniques available for parameter estimation are essentially of two main categories. The first consists of statistical (and sometimes tedious) procedures that have been prescribed for specific forms of reaction kinetics, including generalized mass action (GMA) [16–18], linear-logarithmic (Linlog) [19, 20], and convenience kinetics (CK) [21]. However, it is unclear what technique would then be easily applicable for the myriad combinations of such formulations and new ones that could be prescribed for the individual reactions. Certainly, costly ad hoc experiments [22] or deep learning methods [23–25] could be specified in specific circumstances but a standardized computational procedure would make the learning process more efficient and allows for automation. Thankfully, the predicament can be bypassed by rephrasing the problem generically as one of optimizing a nonlinear objective function in bounded parameter search space (nonlinear programming) [26]. What is only needed in this regard is an optimization algorithm that can traverse the landscape efficiently for sampling the objective function value. The rephrased problem thus trades the ill-defined task of prescribing parameter estimation procedure, by rate law formulation, with a more explicit one of finding an effective optimization algorithm.

The new task is, however, no walk in the park. Ideally, the algorithm must be capable of both exploring and exploiting diverse landscapes to find the global optimal among numerous local solutions. The biological landscape is characterized by terrains more than just numerous and steep optima. It may also be inconsistent because of technical and biological variabilities, and stochasticity, or be distorted by non-representative or inadequate number of datapoints. The terrain may also interchange depending on the specific parameter dimension, and thus exhibits sensitivity to small changes in the search space location. The terrain may further be sensitive to the input data (termed ill-conditioning), due to the prevalence of biochemical feedback/forward loops, for instance, and further aggravated by variability in experimental measurements. For these reasons, the optimization algorithm must minimally be robust, and self-adapt quickly to the local terrain in carrying out its task.

Historically, researchers have recognized the advantage of using stochastic algorithms [9] over deterministic ones (e.g., gradient-based [27] and direct search methods [28, 29]) for the search and optimization of a multi-modal landscape. Among the more effective stochastic methods, evolutionary strategies (ESs) have been consistently reported to be more robust and efficient [30–34] than either simulated annealing (SA) [35] or genetic algorithms (GA) [36]. This may particularly be due to their capacity for self-adapting their strategy parameters, while having all the properties necessary for global optimization [32]. Their outperformance in problems having continuous search space has also been attributed to their specific design for them [26]. On the other hand, SA and GA are originally intended to solve combinatorial problems based on discrete variables [37, 38] with a simplified search space. Indeed, the efficacy of ESs has caught the attention of systems biologists. For example, they have been used as to the explanative modelling of circadian clock [39], development and pattern formation [40–43], and iron metabolism [44]. More broadly, they have been employed to help discern network mechanisms [45], reverse engineer pathways [46, 47], and infer signaling dynamics [48]. In all, the specific application of ESs has been largely based on earlier referenced works and/or through trials and errors. While such qualitative objectives can be adequately met by the ad hoc selection of optimization algorithm, the aspiration of predictive modelling will require a more thorough characterization of promising algorithms in their ability to recover the underlying reaction parameters. For many real world applications, it is as critical that the predictions are made on time [7], thereby placing a premium on the efficiency of the algorithm.

Using both criteria, we ask in this study if there is any evolutionary algorithm (EA) that can efficiently transverse the landscape of major reaction kinetics to find the globally optimal solution. Given the hard nature of the question, we also investigate if there is any value-add in prescribing specific algorithm for the individual formulations of reaction kinetics. In doing so, we recognize there could be no algorithm that outperforms others under all contexts, based on our metrics (no free lunch theorem [50]). Instead, we are searching for acceptable tradeoffs [51, 52] among our key criteria. To this end, we screen the effectiveness of 5 widely used and available EAs in estimating the kinetic parameters of an artificial *in silico* pathway, according to 4 canonical formulations. To mimic an actual application, we otherwise replicate the structure of the mevalonate pathway for limonene production. As a result, we find specific algorithms to be effective under empirical-like conditions, after applying key techniques and learning points to mitigate the effects of measurement errors. Our work thus allows for the more accurate modelling of biological pathway dynamics, as a step towards realizing the aspiration of predictive modelling in systems biology endeavors.

## Materials and methods

### Generation of synthetic enzyme datasets

An overview of the artificial pathway is provided in Figure 1A. For each of 4 *in silico* experiments, the final level for each of 9 enzyme is chosen from three levels: low, medium, and high, via Latin Hypercube sampling [49]. The Hill function is then used to generate the concentration data *e*_*i*_(*t*) of enzyme *i* at time *t* of the run as followed:

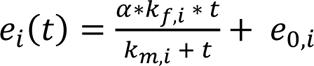

**Figure 1.**
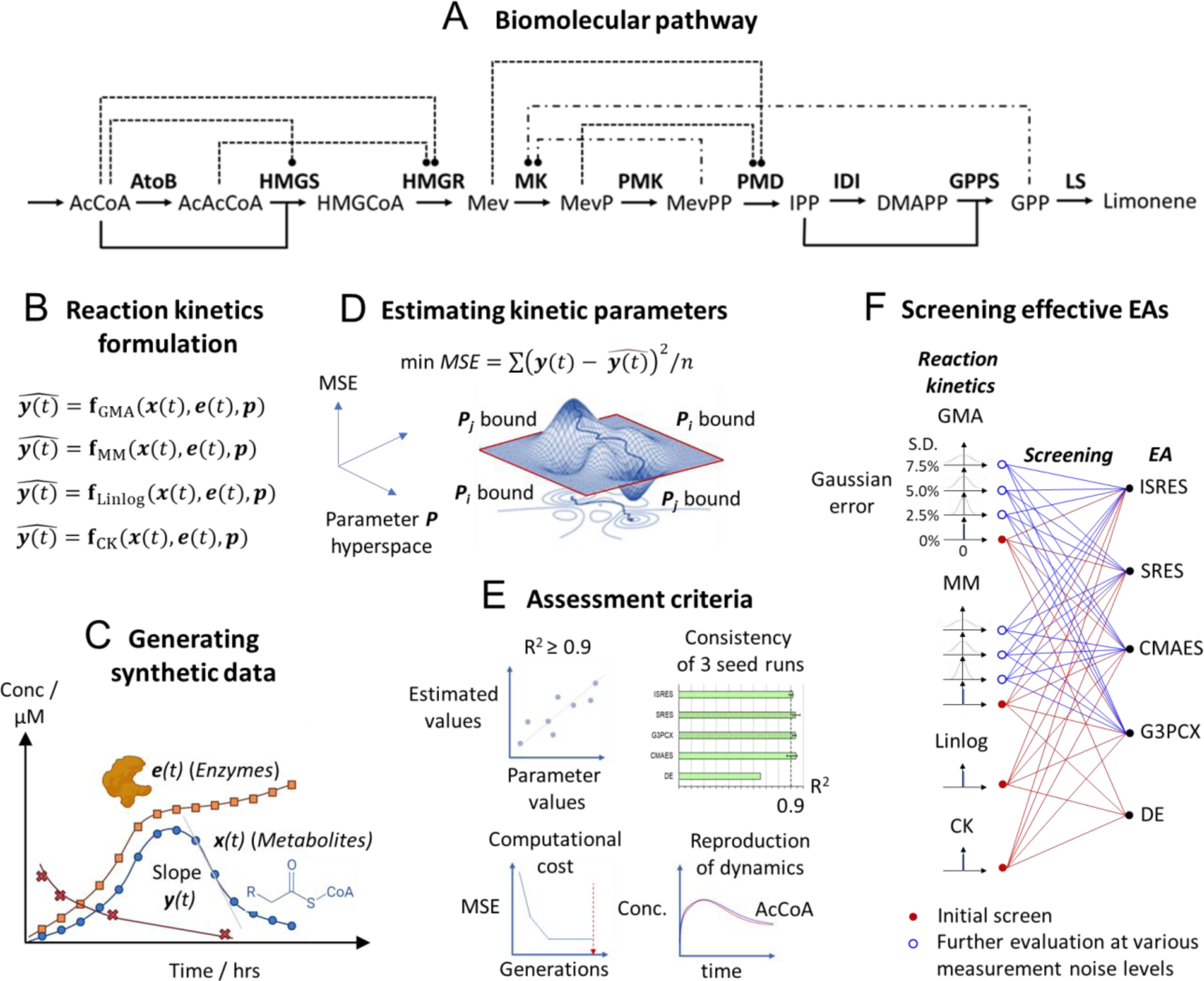
Overview of approach used in study. **A.** Artificial pathway used in study by replicating the topology of the mevalonate pathway for limonene synthesis. Arrows refer to reaction. Dash lines represent feed forward inhibition, whereas mixed dash and dot lines indicate feedback inhibition. **B.** Formulation of separate reaction kinetics for the pathway [generalized mass action (GMA), Michaelis-Menten (MM), linear-logarithmic (Linlog), and convenience kinetics (CK)]. **C.** Time-series data for metabolite concentrations ***x***(*t*) is generated by using the formulated reaction kinetics to simulate reaction progressions through time. The empirical net reaction rate for each metabolite at a given timepoint is then interpreted as the slope of its concentration in time. Separately, the dynamic concentration of enzyme participants ***e***(*t*) is each determined by a specific hyperbolic Hill function. **D.** The mean square error (*MSE*) between empirical net reaction rates and the predicted values based on the kinetic formulation is then minimized as an objective function in kinetic parameter hyperspace. The corresponding parameter values are then reported as an estimation. **E.** The quality of the parameter estimations is then assessed using 4 criteria/considerations: if the coefficient-of-determinant is greater than or equal to 0.9 and is consistently so for three different seed runs, the computational cost (by using the number of generations required for optimization as a proxy), and the degree of reproducibility of the underlying reaction dynamics with the estimated parameters. **F.** Five widely available evolutionary algorithms (EAs) are then screened for their capacity to estimate the parameters of the different kinetic formulations. The initial screening is done using datapoints at 15 minutes interval with no measurement noise. Further evaluations are conducted for selected EAs in estimating GMA and MM parameters at increasing measurement noise and datapoint spacing. The effect of taking parameter average based on different seed solutions as well as data augmentation is also evaluated. **<u>Metabolites</u> AcCoA**: acetyl-coenzyme A; **AcAcCoA**: acetoacetyl-coenzyme A; **HMGCoA**: 3-hydroxy-3-methylglutaryl-coenzyme A; **Mev**: mevalonate; **MevP**: phosphomevalonate; **MevPP**: diphosphomevalonate; **IPP**: isopentenyl pyrophosphate; **DMAPP**: dimethylallyl pyrophosphate; **GPP**: geranyl pyrophosphate **<u>Enzymatic reactions</u> AtoB**: acetoacetyl-CoA thiolase; **HMGS**: hydroxymethylglutaryl-CoA Synthase, **HMGR**: 3-hydroxy-3-methyl-glutaryl-coenzyme A reductase; **MK**: mevalonate kinase; **PMK**: phosphomevalonate kinase; **PMD**: phosphomevalonate decarboxylase; **IDI**: isopentenyl-pyrophosphate delta-isomerase; **GPPS**: geranyl pyrophosphate synthase; **LS**: limonene synthase

Here, *k*_*f*,*i*_ is the maximum increase in enzyme concentration, while the relative amplification factor α is 0.432, 1, and 2.486 for the low, medium, and high final level [50], respectively. Also, *k*_*m*,*i*_is the time required to increase the enzyme concentration by 50% of *α* ∗ *k*_*f*,*i*_, while *e*_0,*i*_is the basal enzyme concentration prior to tuning. Other than α, the parameter values of each enzyme (Data S1) are uniformly sampled from predefined ranges.

### Generation of synthetic metabolite datasets by reaction kinetics

To do so, the initial concentration of metabolites (Data S1) is set so that they are not exhausted during run at the medium tunable level for all enzymes; enzyme concentrations over time are determined by using the Hill’s function as above. Based on these conditions, metabolite concentration data are then generated at 15 minutes interval over 24 hours by simulating the underlying reactions using separate forms of reaction kinetics for the system model (Section one of supplementary information). The kinetic parameter values are obtained as follows: Michaelis-Menten (MM) parameters are first arbitrarily chosen (Data S2) and then used to simulate metabolite time-series data. The latter data is then used to fit the other kinetic models (using CMAES algorithm) to efficiently engender reasonable parameter solutions. Parameters are also liberally altered to keep them within reasonable ranges.

The measurement error of metabolite and enzyme concentrations is sampled from a normal distribution with zero mean and a standard deviation equivalent to 2.5%, 5%, or 7.5% of the underlying value (otherwise referred to as ‘2.5%, 5%, 7.5% noise levels’).

### Computing metabolite net reaction rates from triplicate time-series data

For both metabolite and enzyme datasets, the average of triplicate measurements at regularly spaced matching timepoints is first taken. The net metabolite reaction rate ***y****_t_* at each temporal data point is given by the instantaneous slope, i.e., the first derivative, of the concentration curve in time. We use the Savitzky-Golay filter [51] for evaluation, as it intrinsically smoothens noisy measurements. For datasets with measurement noise, we apply the filter with a temporal window size of 5 datapoints and polynomial order 2. In the absence of noise, the values are 7 and 3, respectively.

To assess the accuracy of the computed ***y*** values, the metabolite concentration at each timepoint t = t_k_ is then recovered by summing up its changes over time from its initial concentration:

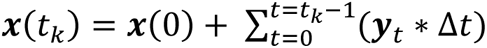

where Δ*t* is the temporal spacing between neighboring datapoints, e.g., Δ*t* = 0.25 hours for 4 temporal datapoints per hour. The regenerated datapoints in time are then compared visually with the corresponding measurements by overlaying them.

### Implementation of evolutionary algorithms, hyperparameters, and termination criteria

We use evolutionary algorithms implemented in the python package, pymoo (version 0.6.0) [14]. Other than setting a population size of 64, the default hyperparameter settings in pymoo are used for all algorithms. We set an upper limit of 1×10^5^ generations for the termination criterion (Table 1), which allows most algorithms to reach a plateau state (in terms of cost function) or to self-terminate for the CMAES algorithm (Figure S1-S4).

**Table 1.**
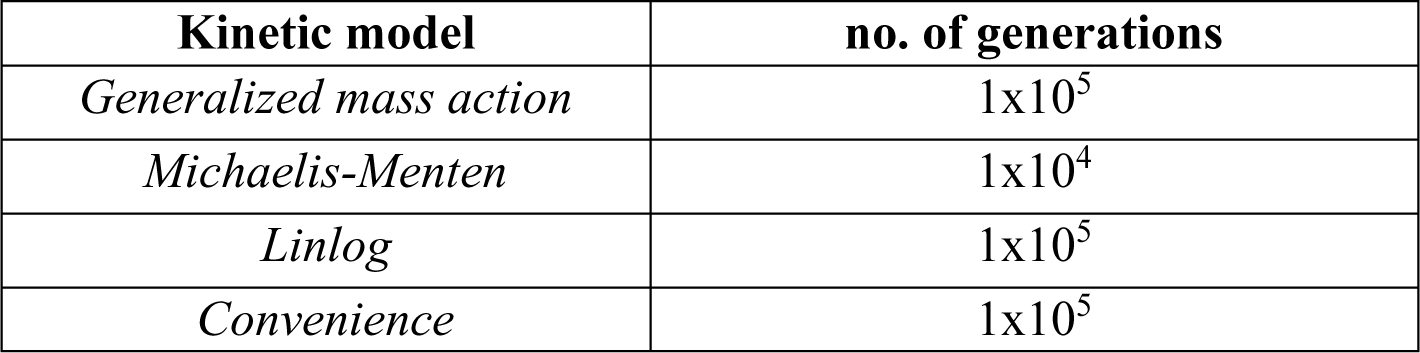
Budgeted number of generations for parameter estimations of various kinetic models.

| **Kinetic model** | **no. of generations** |
| --- | --- |
| *Generalized mass action* | $1 \times 10^5$ |
| *Michaelis-Menten* | $1 \times 10^4$ |
| *Linlog* | $1 \times 10^5$ |
| *Convenience* | $1 \times 10^5$ |

### Estimation of kinetic parameters for artificial pathway

The nonlinear programming problem is formulated as followed:

Find the vector of rate law parameter values (decision variable values), ***p***, to minimize the cost function, i.e., the overall mean square error *MSE* in our case:

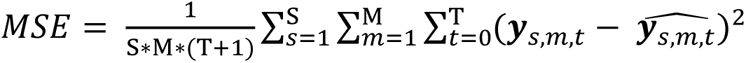

where S is the total number of experimental runs, M is the total number of modelled metabolites, T+1 is the total number of time points, ***y****_s,m,t_* is the net reaction rate for metabolite *m* at time *t* in experiment *s*, and ***ŷ*** is the corresponding prediction by the system model. The model is in the form of a set of ordinary differential equations as described in section one of the supplementary material.

*MSE* is further subjected to:

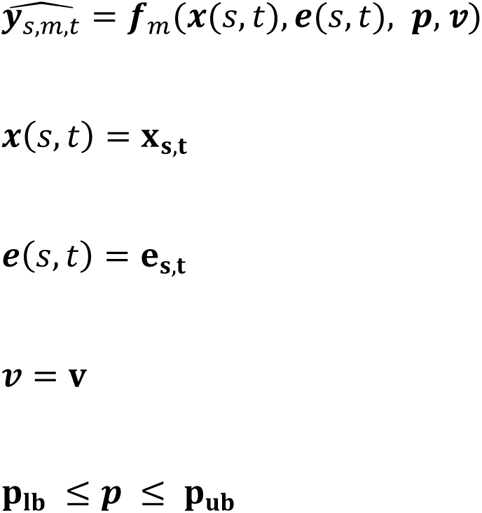

where ***x*** is the vector of metabolite concentrations as state variables, ***e*** is the vector of enzyme concentrations as control variables, **v** is the vector of other parameter values that are not being estimated in the current problem (usually invariant in the system with time). The constant inflow of AcCoA precursor into our modelled system is represented by one such parameter V_in_.

### Performance metric for parameter estimations: coefficient of determination

Recall that the Pearson’s correlation coefficient indicates the degree to which two variables, *y* and *x*, are statistically related by a linear relationship between them, i.e., *y* = m*x* + c. To measure the quality of parameter predictions by candidate evolutionary algorithms, we use a similar metric to gauge the extent to which, the predicted values 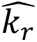 are equivalent to the actual values *k*, i.e.,

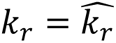

This provides a direct and more stringent measure of estimation quality than otherwise if we merely assess the degree to which *k*_*r*_ and 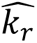 are linearly related. Note that in this case, m and C are 1 and 0, respectively. Analogously, we calculate the coefficient of determinant *R*^2^:

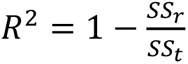

whereby the residual sum of square *SS*_*r*_ and the total sum of square *SS*_*t*_ are also analogously computed as followed:

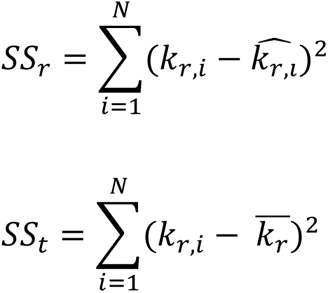

*k*_*r*,*i*_ is the actual value of the *ith* rate law parameter, while 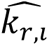 is the corresponding predicted value; 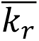 is the mean of the actual value of all N rate law parameters.

## Results and discussion

### Screening evolutionary algorithms for parameter estimations of canonical reaction kinetics

We screen the effectiveness of 5 widely used and available EAs (Overview provided in section two of supplementary information):

i. Differential evolution (DE),
ii. Stochastic Ranking Evolutionary Strategy (SRES),
iii. ‘Improved SRES’ (ISRES),
iv. ‘Covariance Matrix Adaptation Evolution Strategy’ (CMAES), and
v. ‘Generalized generation gap model with parent-centric combination’ (G3PCX)

for 4 kinetic formulations (Section one of supplementary information) of an artificial pathway (Figure 1A and B): GMA [52], MM [53], Linlog [19], and CK [21]. Synthetic data of metabolite dynamics are generated based on each of the reaction kinetics (Figure 1C), preprocessed, and then fitted back to the same formulation (Figure 1D) by minimizing the MSE between predicted and empirical net reaction rates using all candidate EAs.

A total of 4 requirements/considerations are used for evaluating the corresponding parameter solution (Figure 1E):

i. a high quality of parameter estimations, as reflected by the coefficient of determination (R^2^) being greater than 0.9,
ii. consistency in meeting the above requirement based on three different seed solutions,
iii. the number of generations required for the EA to faithfully reach a plateau-state in terms of minimized MSE (figure conservatively rounded up to one significant figure based on the largest of three seeds) and
iv. the ability to reproduce the underlying metabolite dynamics

For criterion iii, we use the number of generations as a convenient proxy for the number of cost function evaluations (which is computationally expensive), as the same population size is used for all algorithms. To discern the effect of kinetic formulation and data quality on the performance of EAs, we first use ‘perfect’ data (Figure 1F), that is, with no measurement noise and a datapoint spacing that is 15 minutes apart. Subsequently, we introduce noise and increase the spacing to ‘strain-test’ the algorithms. This strategy also allows us to efficiently filter out poor performers early on, before progressing to their evaluation under more challenging contexts.

### Evolutionary algorithms differ widely in their effectiveness for parameter estimations

With perfect data, SRES stands out in consistently surpassing our R^2^ threshold for all seed-instances of three out of the four reaction kinetics-of-interest. Its median R^2^ values for GMA, Linlog, and MM kinetics are 0.993, 0.991, and 0.936, respectively (Figure 2, left panel). We note that while SRES was previously reported to produce accurate parameter estimations for a MM model [26], its performance with regard to other reaction kinetics has not been studied.

**Figure 2.**
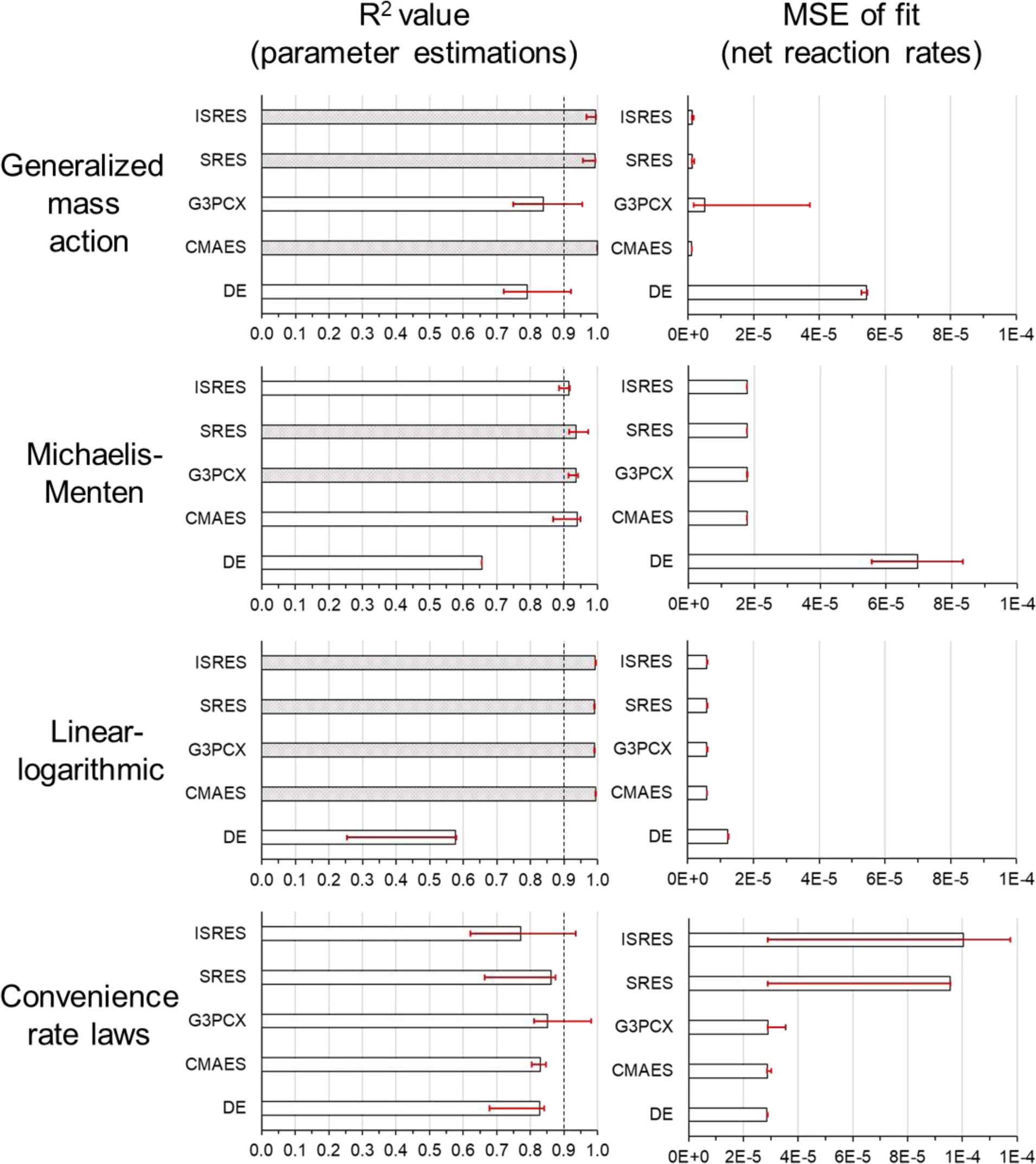
Screening of evolutionary algorithms for parameter estimations of canonical rate law models. Data with no measurement error is used for screening. Both the R^2^-values and the mean square error (MSE) of fit are based on three initial seed solutions. The bars represent the median values while the lower and upper bounds (in red) mark the smallest and largest values based on the seeds, respectively. Note that the computation of the standard deviation as a measure of the spread both above and below the median is inappropriate, as the distribution may not be symmetrical. Vertical dash lines mark the threshold value of 0.9 for R^2^-value metric (R-value of 0.95). A grey bar indicates that the values based on all three seeds have exceeded the threshold. Note that only one R^2^-value has been reported in fitting the Michaelis-Menten model using the differential evolution (DE) algorithm; two others with negative R-value have not been shown.

In the same way, three other EAs exceed the R^2^ threshold for just two classes of reaction kinetics. Specifically, CMAES and ISRES perform similarly well for the same formulations, achieving a respective median R^2^ value of 1.00 and 0.996 for GMA kinetics, and 0.994 and 0.993 likewise for Linlog kinetics. We also find G3PCX to be effective for Linlog formulation (median R^2^ value of 0.99), but it is also as good as SRES in estimating MM parameters (0.936).

Conversely, DE is ineffective for all classes of investigated reaction kinetics, as its R^2^ values mostly fall below 0.9, except for one seed-instance of GMA. Apparently, this is because of the inability of the algorithm to reach a plateau stage in terms of the minimized cost function (MSE-of-fit) (Figure S1-S4), within the budgeted number of generations (Table 1). In contrast, all other EAs can do so for the reaction kinetics that they did well for (as described above). As expected, its MSEs of fit are noticeably higher than those of other EAs for GMA, MM, and Linlog kinetics (figure 2, right panel). Furthermore, in contrast, the other EAs can achieve MSEs of fit within tighter ranges (implying consistency among seed-instances), which are also largely matching with each other (consistency among EAs). For instance, this is true for all other EAs in the case of MM and Linlog kinetics, while (I)SRES and CMAES algorithms are similarly so for GMA kinetics. One learning point from the juxtapositions is that we confirm that the MSE-of-fit, as to its magnitude and consistency among seed-instances, can be helpful in filtering out poor performers. The metric is especially valuable in practical settings, since the R^2^ value cannot be assessed without knowledge of the true solution.

### Effective evolutionary algorithms in the absence of measurement noise

Given the outright poor performance of DE, we dropped it for further evaluation; from this point onward, we use ‘All EAs’ to refer to all other EAs except for DE. We then evaluate the efficacy of the estimated parameters in regenerating metabolite dynamics through simulation. For GMA kinetics, we find CMAES, SRES, and ISRES to closely reproduce all the metabolite dynamics (Figure S5A), which is consistent with their high R^2^ value of parameter estimations (Figure 2, left panel); those of CMAES are in fact virtually indistinguishable from the actual one. Also unsurprisingly given the poor R^2^ performance, G3PCX could not qualitatively reproduce the dynamics of all metabolites for GMA kinetics. For example, we observe a time lag of 4-6 hours for the concentration profile of the 5 most downstream metabolites. However, unexpectedly for MM kinetics, all shortlisted EAs can regenerate the dynamics almost identically (Figure S5B), despite the R^2^ values being appreciably and broadly lower than in the case of GMA kinetics. This suggests the relative insensitivity of MM dynamics to the underlying parameters. We also note the same excellent recovery of Linlog-based dynamics for all EAs (Figure S6), which dovetails with their good metric performance (R^2^ > 0.99). The corresponding MSE-of-fit is also found to be effectively minimized within 5000-50,000 generations (Figure S3, Data S3).

When we further consider computational cost, CMAES stands out in requiring no more than 5000 and 2000 generations for GMA and Linlog kinetics, respectively (Data S3). This corresponds to 1/6 and 1/20 of the generation number required for the next most cost-efficient EA (ISRES: 30,000 and 40,000 respectively) that also meets our other considerations. For MM kinetics, G3PCX is likewise most efficient in requiring only 1000 generations, which is ½ of the other EA (SRES) that fulfils our considerations.

For all that, not a single EA assessed could consistently hit our R^2^ threshold seed-wise for the CK formulation, despite all of them attaining the plateau stage during cost function minimization (Figure S4). The plateaued values further vary noticeably among EAs, suggesting a challenging cost function landscape with numerous local minima. More studies are needed to find an effective optimization method for CK-based models.

### SRES and ISRES are most effective for GMA kinetics in the presence of measurement noise

For the assessment of EAs under experiment-like conditions, we add simulated errors to our metabolite and enzyme ‘measurements’ before repeating our analysis. Again, DE is not considered due to its poor performance in earlier screening. We also exclude Linlog parameter estimations, as the kinetic formulation is a heuristic modelling approach, with no empirical profiling of the rate law to speak of. Given that random errors are expected to increase both the ruggedness and variability of the optimization landscape among replicates, we generate for each initial seed solution a different triplicate dataset to include the effects of measurement noise on parameter predictions. We also take the seed-average of the parameter predictions and compute its R^2^ value to derive a measure of the *expected* quality of parameter estimation.

Contrary to our earlier finding in the absence of noise, CMAES becomes the worst performer for GMA kinetics upon the introduction of noise. Although its expected R^2^ values (reported as filled circles in Figure 3A) and correlation plots at various noise levels (Figure 3B) are not very different from those of other EAs, it clearly exhibits greater variability in the metric value among seed-instances (open circles in Figure 3A), compared to SRES/ISRES algorithms. CMAES likewise exhibits larger variation in the MSE-of-fit compared to other EAs, in the presence of noise (Figure S7, left column). When we test the efficacy of the resulting estimated parameters to recover the mass action dynamics under measurement noise, CMAES is unable to recapture the qualitative profile of the 4 most downstream metabolites, even at a noise level of just 2.5% (Figure S8A) and 5% standard deviation (Figure S8B). At the 7.5% level, the dynamics are off track for 7/10 metabolites (Figure 3C), with an early reaction substrate (AcAcCoA) falsely predicted to be exhausted.

**Figure 3.**
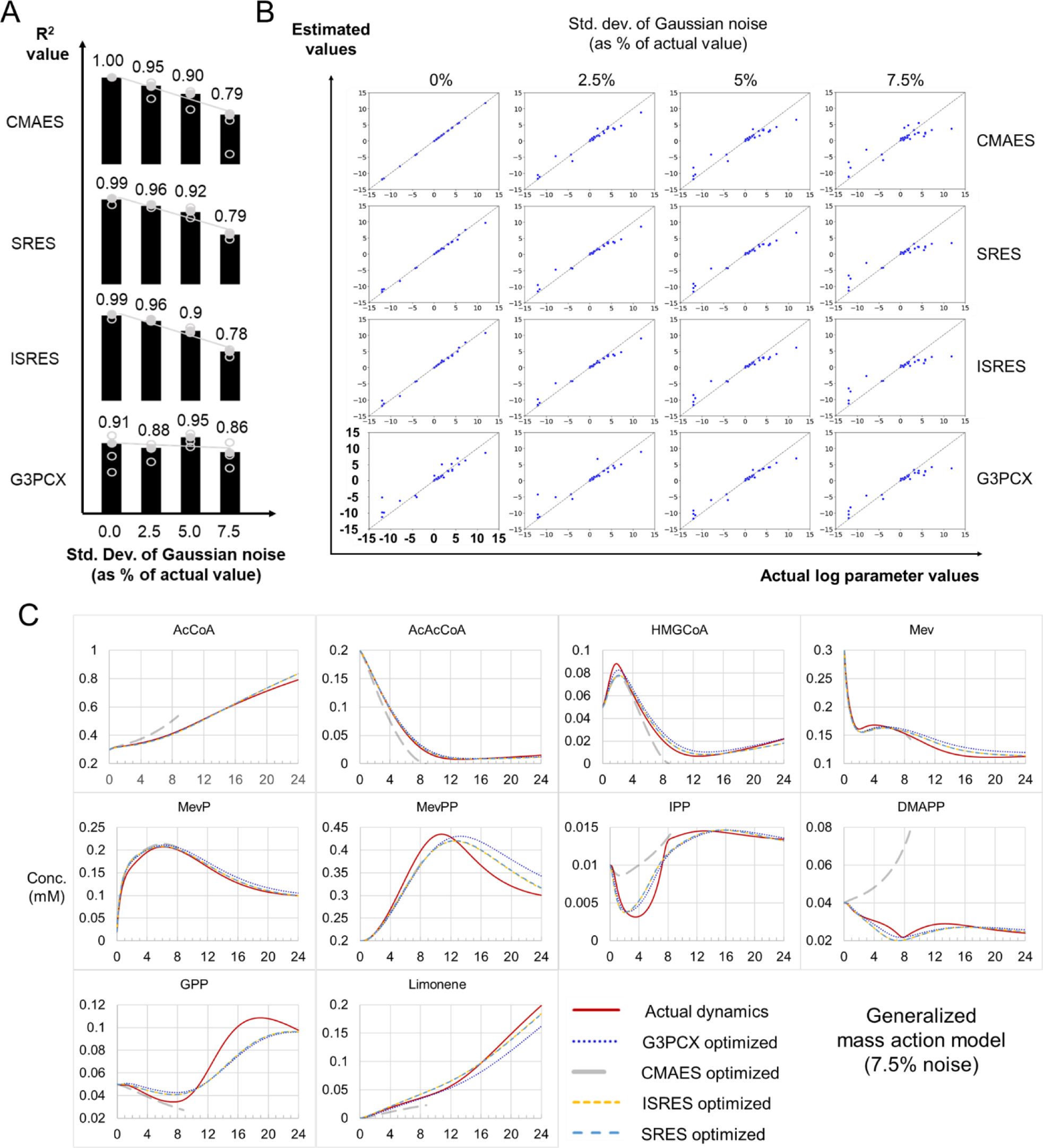
Effectiveness of selected evolutionary algorithms for estimating parameters of generalized mass action model under simulated conditions of increasing measurement errors. **A.** Quality of estimations at increasing measurement errors as quantified by R^2^-value. An open oval represents a value evaluated from a distinct triplicate dataset and random seed solution, whereas a filled oval denotes the R^2^-value of their mean parameter values. The trend of the filled ovals with measurement errors is shown as a straight line. **B.** Correlation plots of estimated parameter values against actual ones at increasing measurement errors for each algorithm. **C.** Simulated metabolite dynamics based on parameters that are estimated using data with measurement errors. Errors are sampled from a normal distribution centered on zero and a standard deviation equivalent to 7.5% of the underlying metabolite or enzyme concentration.

While G3PCX similarly has larger MSE-of-fit and R^2^ variability compared to SRES/ISRES algorithms, this is also the case in the absence of noise. Thus, unsurprisingly, G3PCX cannot recover the qualitative profile of multiple downstream metabolites even with no noise, underscoring its unsuitability for estimating GMA parameters. Instead, SRES/ISRES perform the best for GMA kinetics, generating similar qualitative dynamics at 5% standard deviation or less, albeit at a cost of 100,000 generations.

### With measurement noise, G3PCX remains most effective for recovering MM parameters

We similarly investigate the efficacy of EAs for estimating MM parameters when measurement errors are present. In this regard, the performance of all algorithms based on expected R^2^ value remains relatively stable at increasing measurement noise (Figure 4A), compared to the estimation of GMA parameters. To illustrate, the metric stays between 0.9 and 0.96 (filled ovals) for all EAs even at 7.5% noise level, whereas the same figure ranges between 0.78 and 1 for GMA parameter estimations. To view in another way, there is no discernable drop in the expected R^2^ value for both ISRES and G3PCX algorithms with increasing noise, while those of CMAES and SRES decrease minorly (≤ 0.06). In contrast, we observe a decrease of 0.21 in the metric value for three EAs (CMAES, SRES, ISRES) with increasing noise, for the case of GMA kinetics. The similar correlation plots between the mean estimated parameters and actual values at increasing measurement noise further illuminate the more insensitive nature of MM parameter estimations to measurement errors (Figure 4B).

**Figure 4.**
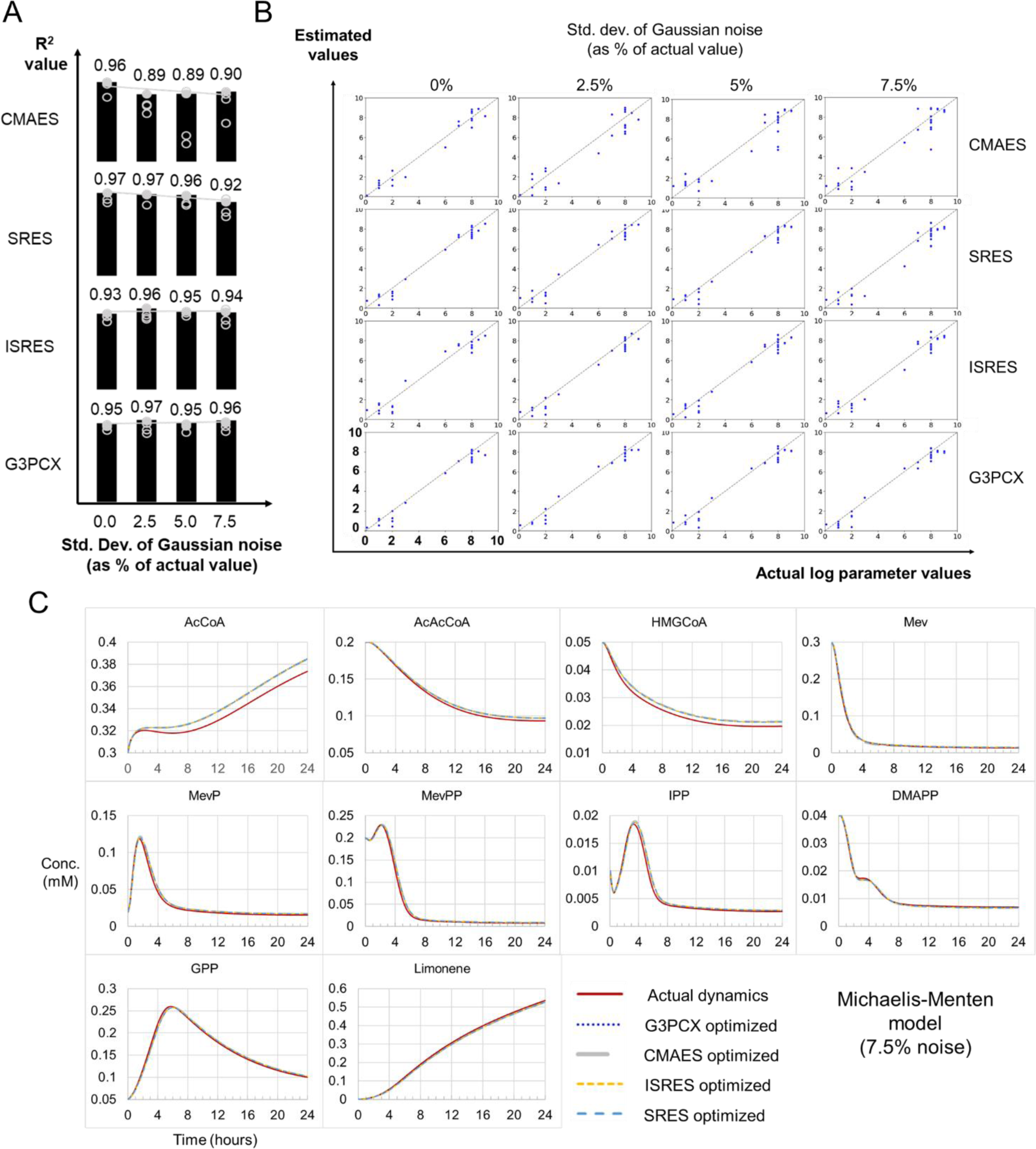
Effectiveness of selected evolutionary algorithms for estimating parameters of Michaelis-Menten model under simulated conditions of increasing measurement noise. Refer to Figure 3 for the legend of corresponding sub-figures **A-C**.

The EAs also recover MM dynamics more reliably than that of GMA dynamics, as demonstrated by the largely unchanging metabolite profiles that are generated for MM kinetics with increasing noise for all EAs. Specifically, the profiles of all EAs are virtually identical to the actual dynamics at 0% (Figure S5B) and 2.5% noise levels (Figure S9A), while showing marginal difference for the three most upstream metabolites at 5% (Figure S9B) and 7.5% noise levels (Figure 4C). In this regard, all EAs perform similarly well in recovering MM parameters and dynamics when measurement noise is present. However, after taking computational cost into account, G3PCX remains the most effective, by necessitating only 2000 generations to do so (Figure S10). The figure is 66% that of the next most efficient EA (SRES: 3000 in Data S3) that satisfactorily meets all our other considerations. It would be interesting to verify for a variety of network topologies, if the measurement noise has, indeed, less effect on recovering the parameters and dynamics of a MM formulation, compared to its GMA equivalence.

### CMAES is most sensitive to measurement noise

In addition to the case for GMA kinetics, we observe CMAES to show increasing inconsistency in MM parameter estimations with more measurement noise, as reflected by the larger variability in R^2^ value (open ovals in Figure 4A). Note that this is to a greater extent than other EAs. Taken together, the findings for both GMA and MM kinetics implicate the unreliability of CMAES under noisy experimental conditions.

We further note the inconsistency in MM parameter estimation by CMAES, as discussed above, has not been obvious from the distribution of the MSE-of-fit (Figure S7, right column). Instead, the MSE value for individual seed-instances is largely identical to that of other EAs, resulting in an akin overall distribution for all noise levels. Clearly, similar MSE values does not necessarily imply similar quality of parameter estimations. (Yet conversely, we reason that MSE values and distributions that are distinctly larger should be seen as strong indicators of the relative ineffectiveness of parameter estimations.) Notably, the sensitivity of CMAES to noisy data would not be obvious, if not for the usage of R^2^ metric. In turn, the usage of the latter metric would only be possible if the underlying parameter values and reaction kinetics are known, such as in the context of an *in silico* pathway.

## Concluding remarks

In pushing the performance of EAs to their limit, we realize the feasibility of deriving remarkably good parameter estimates in the presence of measurement errors, by using the appropriate EA (Table 2) with an entire repertoire of mitigation techniques for our approach (Section three of supplementary information). However, there remain various caveats. For example, some algorithms such as CMAES may be intrinsically unable to adapt to the variability of biological data. Also, the bounds of the parameter search space have enormous effect on whether the true solution can ever be found. While we recommend constraining as much as possible the search space guided by domain knowledge, it may not be practically feasible for most parameters. It is also currently unclear if an EA is intrinsically suitable for the reaction kinetics that it performs well for in our study, and to what extent, it will remain so with increasing number of reactions. Besides the need to address these questions, future works should explore if the hyperparameters tuning of EAs can make a material difference to their performance, such as in the required computational cost. To appreciate the latter’s importance, imagine an early fish requiring minutes just to scan possible escape routes in the face of an ambush predator. Clearly, there is a lot more to learn from the history of life if predictive modelling is to become truly useful for systems biology endeavors.

**Table 2.** Suggested evolutionary algorithms (EAs) for parameter estimations of known underlying reaction kinetics.

| System/model | Suggested EAs | Comment | Supporting figure(s) |
| --- | --- | --- | --- |
| *Generalized mass action system (without measurement error)* | CMAES | Most consistent & highest quality of parameter estimations, lowest computational cost | Figure 2 & S5A, data S3 |
| *Generalized mass action system (with measurement errors)* | (I)SRES | Most consistent & highest quality of parameter estimations | Figure 3 & S8 |
| *Michaelis-Menten system (with/without measurement error)* | G3PCX | Lowest computational cost, among the best in consistency & quality of parameter estimations | Data S3, figure 2, 3, S5B, S9 |
| *Linear logarithmic model<sup>^</sup>* | CMAES | Lowest computational cost, among the best in consistency & quality of parameter estimations | Figure 2, data S3 |

<sup>^</sup> The kinetic formulation is a heuristic modelling approach, with no empirical profiling of the rate law to speak of.

## Supporting information

Synthetic omics data

## Key Points

- Systems biology has begun its nascent transition from explanatory modelling of fitted data to the predictive modelling of unseen data. However, it still necessitates the effective recovery of all underlying reaction parameters.
- We found algorithms that are effective under marked measurement noise for specific reaction kinetics, as a step towards predictive modelling for systems biology endeavors.
- We further prescribe the key learning points and approach to do so.

## Acknowledgments

HCY conceptualized, planned, and conducted the research. HCY supervised student and wrote the article. VV explored various parameter combinations for the Savitzky-Golay filter, and methods for pooling solutions based on multiple seeds. KS supervised the work and reviewed the article. Publicly available codes were modified or reused for our work [8].

## Data availability

The synthetic omics data underlying this article are available as online supplementary material. Data regarding optimization runs will be shared on reasonable request to the corresponding author.

## Abbreviations

GMA: generalized mass action
MM: Michaelis-Menten
Linlog: linear-logarithmic
CK: convenience kinetics

