## Supplementary information for "Identifying effective evolutionary strategies for uncovering reaction kinetic parameters under the effect of measurement noises"

#### 1. System model

$\hat{\boldsymbol{y}}$ , the vector of model prediction for the net reaction rate of individual metabolites can be represented as followed:

$$\hat{\boldsymbol{y}} = \begin{bmatrix} \frac{d[AcCoA]}{dt} \\ \frac{d[AcAcCoA]}{dt} \\ \frac{d[HMGCoA]}{dt} \\ \frac{d[Mev]}{dt} \\ \frac{d[MevP]}{dt} \\ \frac{d[MevPP]}{dt} \\ \frac{d[IPP]}{dt} \\ \frac{d[DMAPP]}{dt} \\ \frac{d[GPP]}{dt} \\ \frac{d[Limonene]}{dt} \end{bmatrix}$$

whereby each vector element (net rate of reaction of a metabolite) can be evaluated by summing the contributions from individual reactions (Figure 1 in main paper):

$$\frac{d[AcCoA]}{dt} = r_{v_{in}} - r_{AtoB} - r_{HMGS}$$

$$\frac{d[AcAcCoA]}{dt} = r_{AtoB} - r_{HMGS}$$

$$\frac{d[HMGCoA]}{dt} = r_{HMGS} - r_{HMGR}$$

$$\frac{d[Mev]}{dt} = r_{HMGR} - r_{MK}$$

$$\frac{d[MevP]}{dt} = r_{MK} - r_{PMK}$$

$$\frac{d[MevPP]}{dt} = r_{PMK} - r_{PMD}$$

$$\frac{d[IPP]}{dt} = r_{PMD} - r_{IDI} - r_{GPPS}$$

$$\frac{d[DMAPP]}{dt} = r_{IDI} - r_{GPPS}$$

$$\frac{d[GPP]}{dt} = r_{GPPS} - r_{LS}$$

$$\frac{d[Limonene]}{dt} = r_{LS}$$

The rate of the individual reactions is as followed, based on:

1. Generalized mass action kinetics

$r_{AtoB} = 10^{k_{11}}[AtoB]^{k_{12}}[AcCoA]^{k_{13}}$

$r_{HMGS} = 10^{K_{21}}[HMGS]^{K_{22}}[AcCoA]^{k_{23}}[AcAcCoA]^{k_{24}}$

$r_{HMGR} = 10^{K_{31}}[HMGR]^{K_{32}}[AcCoA]^{-k_{33}}[AcAcCoA]^{-k_{34}}[HMGCoA]^{k_{35}}$

$r_{MK} = 10^{K_{41}}[MK]^{K_{42}}[Mev]^{k_{43}}[GPP]^{-k_{44}}[MevPP]^{-k_{45}}$

$r_{PMK} = 10^{K_{51}}[PMK]^{K_{52}}[MevP]^{k_{53}}$

$r_{PMD} = 10^{K_{61}}[PMD]^{K_{62}}[MevPP]^{k_{63}}[MevP]^{-k_{64}}[Mev]^{-k_{65}}$

$r_{IDI} = 10^{K_{71}}[IDI]^{K_{72}}[IPP]^{k_{73}}$

$r_{GPPS} = 10^{K_{81}}[GPPS]^{K_{82}}[IPP]^{k_{83}}[DMAPP]^{k_{84}}$

$r_{LS} = 10^{K_{91}}[LS]^{K_{92}}[GPP]^{k_{93}}$

$r_{v_{in}} = 10^{v_{in}}$

2. Michaelis-Menten kinetics

The rate laws are taken from the work of *Costello and Martin* [1]. For $r_{Mk}$, the following is quoted from their supplementary material, which indicates that *MevPP*, and not *MevP* is an inhibitor of the reaction [2]:

*'…diphosphomevalonate (DPM) is an noncompetitive inhibitor with respect to both substrates. DPM binds at an allosteric site, and inhibition cannot be overcome by an increasing substrate concentration…'*

Accordingly, we correct the typo in our model.

$$r_{AtoB} = \frac{10^{k_{11}}[AtoB][AcCoA]}{10^{k_{12}} + 10^{k_{13}}[AcCoA]}$$

$$r_{HMGS} = \frac{10^{k_{21}}[HMGS][AcCoA][AcAcCoA]10^{k_{s3}}}{10^{k_{22}}[AcAcCoA] + 10^{k_{23}}[AcCoA] + 10^{k_{24}}[AcCoA][AcAcCoA]}$$

$$r_{HMGR} = \frac{10^{k_{31}}[HMGR][HMGCoA]}{10^{k_{32}}[AcCoA] + 10^{k_{33}}[AcAcCoA] + 10^{k_{34}}[HMGCoA] + 10^{k_{35}}}$$

$$r_{Mk} = \frac{10^{k_{41}}[MK][Mev]}{10^{k_{42}}[GPP] + 10^{k_{43}}[MevPP] + 10^{k_{44}}[Mev] + 10^{k_{45}}}$$

$$r_{PMK} = \frac{10^{k_{51}}[PMK][MevP]}{10^{k_{52}} + [MevP]}$$

$$r_{PMD} = \frac{10^{k_{61}}[PMD][MevPP]}{10^{k_{62}}[MevP] + 10^{k_{63}}[Mev] + 10^{k_{64}}[MevPP] + 10^{k_{65}}}$$

$$r_{IDI} = \frac{10^{k_{71}}[IDI][IPP]}{10^{k_{72}} + [IPP]}$$

$$r_{GPPS} = \frac{10^{k_{81}}[GPPS][IPP][DMAPP]}{10^{k_{82}} + 10^{k_{83}}[IPP] + 10^{k_{84}}[DMAPP] + [IPP][DMAPP]}$$

$$r_{LS} = \frac{10^{k_{91}}[LS][GPP]}{10^{k_{92}} + [GPP]}$$

$$r_{v_{in}} = 10^{v_{in}}$$

#### 3. Linear logarithmic kinetics

$$r_{AtoB} = 10^{k_{11}}[AtoB] + 10^{k_{22}}log_{10}[AcCoA]$$

$$r_{HMGS} = 10^{k_{21}}[HMGS] + 10^{k_{22}}log_{10}[AcCoA] + 10^{k_{23}}log_{10}[AcAcCoA]$$

$$r_{HMGR} = 10^{k_{31}}[HMGR] - 10^{k_{32}}log_{10}[AcCoA] - 10^{k_{33}}log_{10}[AcAcCoA] + 10^{k_{34}}log_{10}[HMGCoA]$$

$$r_{MK} = 10^{k_{41}}[MK] + 10^{k_{42}}log_{10}[Mev] - 10^{k_{43}}log_{10}[GPP] - 10^{k_{44}}log_{10}[MevPP]$$

$$r_{PMK} = 10^{k_{51}}[PMK] + 10^{k_{52}}log_{10}[MevP]$$

$$r_{PMD} = 10^{k_{61}}[PMD] + 10^{k_{62}}log_{10}[MevPP] - 10^{k_{63}}log_{10}[MevP] - 10^{k_{64}}log_{10}[Mev]$$

$$r_{IDI} = 10^{k_{71}}[IDI] + 10^{k_{72}}log_{10}[IPP]$$

$$r_{GPPS} = 10^{k_{81}}[GPPS] + 10^{k_{82}}log_{10}[IPP] + 10^{k_{83}}log_{10}[DMAPP]$$

$$r_{LS} = 10^{k_{91}}[LS] + 10^{k_{92}}log_{10}[GPP]$$

$$r_{v_{in}} = 10^{v_{in}}$$

#### 4. Convenience kinetics

$$r_{AtoB} = 10^{kcat_1}[AtoB]\frac{\widehat{s_{11}}^2}{1+\widehat{s_{11}}+\widehat{s_{11}}^2}, where\ \widehat{s_{11}} = \frac{[AcCoA]}{10^{kma_{11}}}$$

$$r_{HMGS} = 10^{kcat_2}[HMGS]\frac{\widehat{s_{21}}\widehat{s_{22}}}{1+\widehat{s_{21}}+\widehat{s_{22}}+\widehat{s_{21}}\widehat{s_{22}}}, where\ \widehat{s_{21}} = \frac{[AcCoA]}{10^{kma_{21}}}, \widehat{s_{22}} = \frac{[AcAcCoA]}{10^{kma_{22}}}$$

$$r_{HMGR} = 10^{kcat_3}h_{31}h_{32}[HMGR]\frac{\widehat{s_{33}}}{1+\widehat{s_{33}}}, where\ h_{31} = \frac{10^{KI_{31}}}{10^{KI_{31}}+[AcCoA]}, h_{32} = \frac{10^{KI_{32}}}{10^{KI_{32}}+[AcAcCoA]}, \widehat{s_{33}} = \frac{[HMGCoA]}{10^{kma_{33}}}$$

$$r_{MK} = 10^{kcat_4}h_{46}h_{49}[MK]\frac{\widehat{s_{44}}}{1+\widehat{s_{44}}}, where\ h_{46} = \frac{10^{KI_{46}}}{10^{KI_{46}}+[MevPP]}, h_{49} = \frac{10^{KI_{49}}}{10^{KI_{49}}+[GPP]}, \widehat{s_{44}} = \frac{[Mev]}{10^{kma_{44}}}$$

$$r_{PMK} = 10^{kcat_5}[PMK]\frac{\widehat{s_{55}}}{1+\widehat{s_{55}}}, where\ \widehat{s_{55}} = \frac{[MevP]}{10^{kma_{55}}}$$

$$r_{PMD} = 10^{kcat_6}h_{64}h_{65}[PMD]\frac{\widehat{s_{66}}}{1+\widehat{s_{66}}}, where\ h_{64} = \frac{10^{KI_{64}}}{10^{KI_{64}}+[Mev]}, h_{65} = \frac{10^{KI_{65}}}{10^{KI_{65}}+[MevP]}, \widehat{s_{66}} = \frac{[MevPP]}{10^{kma_{66}}}$$

$$r_{IDI} = 10^{kcat_7}[IDI]\frac{\widehat{s_{77}}}{1+\widehat{s_{77}}}, where\ \widehat{s_{77}} = \frac{[IPP]}{10^{kma_{77}}}$$

$$r_{GPPS} = 10^{kcat_8}[GPPS]\frac{\widehat{s_{87}}\widehat{s_{88}}}{1+\widehat{s_{87}}+\widehat{s_{88}}+\widehat{s_{87}}\widehat{s_{88}}}, where\ \widehat{s_{87}} = \frac{[IPP]}{10^{kma_{87}}}, \widehat{s_{88}} = \frac{[DMAPP]}{10^{kma_{88}}}$$

$$r_{LS} = 10^{kcat_9}[LS]\frac{\widehat{s_{99}}}{1+\widehat{s_{99}}}, where\ \widehat{s_{99}} = \frac{[GPP]}{10^{kma_{99}}}$$

$$r_{v_{in}} = 10^{v_{in}}$$

#### 2. Candidate evolutionary algorithms

The following candidates are evaluated in our work. Note that clustering techniques [3, 4] and adaptive stochastic methods [5, 6] are not considered, as they are deemed less effective [7]. Ant Colony optimization, Taboo Search, and particle swarm methods [8] are not commonly used, and thus also not assessed.

**1. Differential evolution** (DE) [9] is a direct global search method, which is designed for optimizing nonlinear, non-differentiable, continuous objective function. It iteratively improves a population of candidate vector solutions as followed:

- Initialization A population of candidate solutions is first initialized.
- Mutation For each candidate solution, three random individuals (a, b, c) are selected from the population and combined to form a trial vector by using the operation: trial_vector = a + F * (b - c), where F is a user-defined amplification factor of (b–c).
- Crossover operation then results in a segment of the trial vector replacing its counterpart in a predetermined target. The starting location of the segment is randomly chosen, while a user-defined parameter, "crossover probability" (CR), determines the probability of extending the segment by an additional vector component.
- Selection The resulting trial vector is then compared to the target based on objective function value and replaces the target if it gives a better value.
- Termination The mutation, crossover, and selection steps are repeated until a termination condition is met, such as a predefined number of generations, convergence of the objective function to some plateau value, etc.

**2. Evolutionary strategies** (ESs) [10] are population based like DE, but are designed for search in continuous space. They are highly varied in the specifics of implementation but generally consist of the following stages:

- Initialization: μ number of parental vector solutions are first initialized, based on user-defined sampling scheme.
- Recombination: the μ parental solutions is duplicated, and then followed by the probabilistic exchange of solution components among them, via predefined rules, to generate a new population of λ 'offspring' solutions. Note that the process is carried out for the CMAES algorithm, but not SRES and ISRES.
- Mutation: the change in each component value of the 'offspring' solutions is sampled from the corresponding normal distribution with zero mean and some standard deviation. The latter, also called the mutation step size, is a search strategy parameter, as it has the effect of balancing the exploration and exploitation of solutions. For example, the mutation step size can be increased to ensure a wider search when progress is poor, and to decrease the step size when progress is promising. ES variants may implement specific approach to automatically adapt the strategy parameter to the search landscape. Offspring solutions generated during the process may only be accepted ('non-lethal') if they are within user-defined bounds.
- Selection: the fitness of each of λ offspring is then evaluated based on the objective function, with the best μ solutions chosen to make up the next generation of parents. The μ parents of the current generation may form part of the selection pool (not the case for our tested variants).
- Termination: the recombination, mutation, and selection steps are repeated until a termination condition is met, such as a predefined number of generations, convergence of the objective function to some plateau value, etc.

In our study, three ES variants were tested:

**2.1 Stochastic Ranking Evolutionary Strategy** (SRES) [11] probabilistically uses only objective function for comparing adjacently ranked solutions during bubble sort, without penalizing infeasible solution(s). It thus gives unacceptable solutions a chance to be selected, which could give rise to better and feasible solutions in the next generation. In this way, the algorithm encourages exploration and help prevent premature convergence.

**2.2 'Improved SRES'** (ISRES) attempts to improvise over SRES by incorporating an alternative mutation strategy of using the differential between selected solutions to generate new ones [12], similarly to DE. This is carried out for variables in the search space that are dependent on others.

**2.3 'Covariance Matrix Adaptation Evolution Strategy'** (CMAES) [13] adapts the multi-variate normal distribution that is used for sampling mutation sizes, by overweighting those selected for previously. As such, some solution components may have similar, and thus correlated, preferred mutation sizes. In this light, the covariance matrix of the distribution may become non-diagonal over generations. Besides adapting the covariance matrix, the algorithm also adopts specific principles used in natural gradient descent to ensure that the distribution remains undistorted by one of its key step (reparameterization).

**3. 'Generalized generation gap model (G3P) with parent-centric combination'** (G3PCX)[14] presumes offspring solutions generated near parental ones in continuous search space to be more likely to be good candidates, given that the parents themselves have been previously selected for based on their fitness. A particular recombination operation is used to ensure the offspring solutions are normally distributed and centred on individual parents (‘parent-centric’). The algorithm also modifies its core model, the minimal generation gap (MGG) [15], to select the best two solutions separately from each batch of similar fitness in each generation, instead of using the roulette-wheel selection procedure, which is computationally expensive, to choose just one solution.

### 3. Learning points and approach for effective estimation of kinetic parameters

#### 3.1 Taking average from multiple seed solutions improves accuracy and reliability of estimations

From Figure 3A and 4A, we also newly observe the averages of parameter values from different seed solutions to have $R^2$ values that consistently surpass or are in the top range of the individual values (respective filled circles versus open circles in the figures). This result is regardless of the tested optimization algorithms, the measurement noise levels, and the reaction kinetics being fitted. Motivated by the potentially useful finding, we investigate if this remains true with increasing number of seed solutions, and if so, whether the resulting $R^2$ does converge to some significantly higher value for the purpose of practical application. To do so, we use the same triplicate datasets for 100 seeds to investigate the sole effect of their increasing number.

For the inquiry, we use G3PCX optimization of the MM formulation at '5% measurement noise' for the data, due to its practical relevance and fast turnaround. As depicted in Figure S11, we find the beneficial effect of taking average to be quickly evident for small cumulative seed numbers and to further increase and rapidly stabilize at a higher $R^2$ value of 0.97 beyond 20-40 seeds. In contrast, the global average value is 0.92, indicating an appreciable enhancement of accuracy by 0.05 in terms of the expected $R^2$ value (P-value of Wilcoxon-Mann-Whitney test [16] = $2.93 \times 10^{-42}$ for 100 seeds, $3.73 \times 10^{-4}$ for 10 seeds). In addition, while the $R^2$ value fluctuates greatly among individual runs (0.69 to 0.98), the same metric based on average parameter values is more reliable, remaining steady at 0.97. A similarly effective outcome can also be achieved by taking the median of parameter solutions (Figure S11, P-value of Wilcoxon-Mann-Whitney test = $3.66 \times 10^{-43}$ for 100 seeds, $4.65 \times 10^{-4}$ for 10 seeds). Indeed also, the averages/medians of parameter values from different seed solutions do have $R^2$ values, which consistently surpass or are in the top range of the individual values. We thus recommend taking the average/median of parameter values based on multiple seed solutions to arrive at a more accurate and reliable set of values than would be expected individually.

**3.2 Parameters and dynamics recovery can benefit from datapoints as close as 15 minutes apart**

Given the superior applicability of SRES and G3PCX algorithms to the respective case of GMA and MM kinetics under noisy empirical conditions, we further investigate their sensitivity to another common factor affecting performance: the number of hourly empirical datapoints. To do so, we reduce the number of datapoints from four per hour to (i) two, and then (ii) one hourly. Then, in each case, we reevaluate the capability of the two EAs regarding parameter estimations and the recovery of metabolite dynamics.

For systems having reversals in dynamic trend (i.e., switching from increasing to decreasing concentrations, and vice versa) between two to four hours like our modelled system, our findings suggest the benefits of having measurements more than once or even twice hourly. For a start, there is a tendency towards more accurate parameter estimations as exemplified by the $R^2$ value for the SRES case study, regardless of measurement noise (Figure S12, top and bottom of left column). (Note that there is no clear deterioration in the recovered dynamics due to less datapoints [Figure S13 versus Figure S5A and S8B]). Although the G3PCX estimation of MM parameters shows similarly high and stable metric value for all considered cases of hourly datapoints (one, two, and four) in the absence of measurement noise (Figure S12, top and bottom of right column), the algorithm however performs poorly according to the metric for the case of one hourly datapoint with measurement noise. More illuminatingly, when we simulate the MM dynamics with the estimated parameters, we find the profiles based on one or two hourly datapoints without measurement noise do not faithfully reproduce all the underlying profiles (Figure S14), unlike the case when four hourly datapoints are used (Figure S5B). On the same note, in the presence of measurement noise, the regenerated dynamics of two hourly datapoints (Figure S14) are not as close to the real profile, compared to 4 datapoints (Figure 4B). Taken together, our data suggests that having more than two hourly empirical datapoints (e.g., four) can be still helpful for improving parameter estimations and the recovery of dynamics, regardless of noise; certainly, this is of paramount importance in high value applications, and when safety is critical [17].

##### 3.3 No evidence that data augmentation improves parameter estimations

Our finding in the previous section begs the question if the augmentation of dynamic data could improve the recovery of parameters and metabolite dynamics, although it has been clear to us that it cannot be a substitute for empirical data. Again, we use the case studies of SRES and G3PCX algorithms for GMA and MM kinetics, respectively, both in the presence and absence of measurement noise. Initial datasets with one or two hourly datapoints are then augmented to four datapoints hourly before parameter estimations. We find, at best, a small and negligible increase in the $R^2$ value of parameter estimations in many cases (Figure S15) but can also potentially result in a sharp and significant decrease in the accuracy of estimations, as revealed by the data augmentation of one hourly datapoint for the G3PCX estimation of MM parameters (Figure S15, bottom right). As such, we refrain from recommending data augmentation for the purpose of parameter estimation, as there is no evidence of its benefit from our analysis. In fact, our finding of a clearly adverse effect in one case suggests that the repeated usage of data augmentation could result in a poorer quality of parameter estimations on average.

##### 3.4 Two key modules for parameter estimation

To facilitate the application of our learning points and approach for future works, we recapitulate the two key modules and component steps contributing to effective parameter estimations (Figure S16). Module one (first 5 steps in red) focuses on deriving high quality estimates of the net reaction rate for all metabolites at each timepoint. In doing so, it forms the foundation for accurate parameter estimations in module two (last three steps in blue). Their rationale or learning points are briefly as follows:

**Module one**

- In step one, the generation of timepoint data that are as close as possible reflects the value of having more timepoints in deriving accurate empirical values of net reaction rates. We exemplify this point by demonstrating the noticeable end improvement in parameter estimation accuracy by simply increasing from one to two and then four hourly datapoints (Figure S12). The effect is even more pronounced in the typical case where measurement noise is present. This is despite the implementation of all the other steps for maximizing accuracy. Note that data augmentation is not an effective solution for insufficient timepoints (Figure S15).
- Steps two and three include means for diminishing the effect of noise: taking average of triplicate measurements and employing the Savitzky-Golay filter, which concurrently filters out noise, while computing the net reaction rate (materials and methods).
- Steps 3-5 further provide the mechanism for generating realistic time profiles of metabolites based on net reaction rates, by adjusting the Savitzky-Golay parameters (window size and polynomial degree for local fitting). By observing the similarity with experimental profiles, we obtain direct evidence of the reliability of derived net reaction

rates. Among other possibilities, the MSE between regenerated and empirical concentration values (the average of triplicates) across all timepoints and metabolites can be used as a metric for assessing their similarity in an automated search for good Savitzky-Golay parameters.

**Module two**

- Step 6 estimates kinetic parameters (materials and methods) by using effective EA implementation according to the underlying rate law (Table 2), and their corresponding hyperparameters and termination criteria (materials and methods).
  Importantly, the average of parameter values from multiple seed solutions (e.g., ≥10) is taken to improve their accuracy and decrease variability, compared to individual seed runs (Figure S11).
- Steps 6-8 are collective mechanism for enhancing the similarity between regenerated and experimental time profiles of metabolites, like steps 3-5. Without knowing the actual kinetic parameter values, the similarity serves as the end-all-be-all criteria for assessing the overall reliability of their estimations. The bounds of the parameter search space should be informed by domain knowledge as much as possible. The hyperparameters of the employed EA can also be tuned.

By deconstructing the approach into two modules, it will be also easier to sieve out issues as to data quality and preprocessing, and the actual parameter estimations, by examining the quality of regenerated profiles in module one and two, respectively.

#### 4. Supplementary Figures

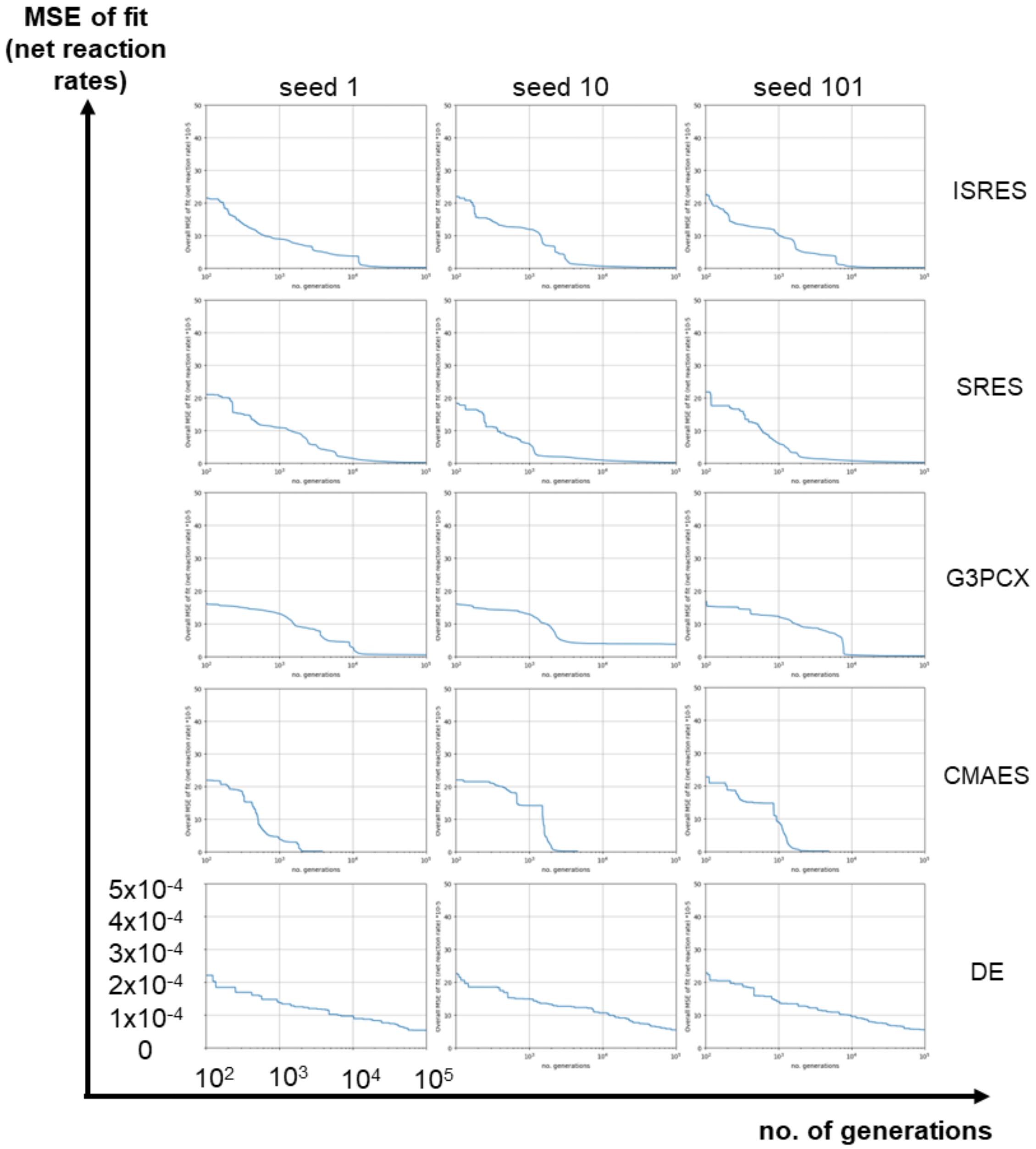


**Figure S1. Convergence profile in fitting the *generalized mass action* model by candidate evolutionary algorithms and seed-instances.**

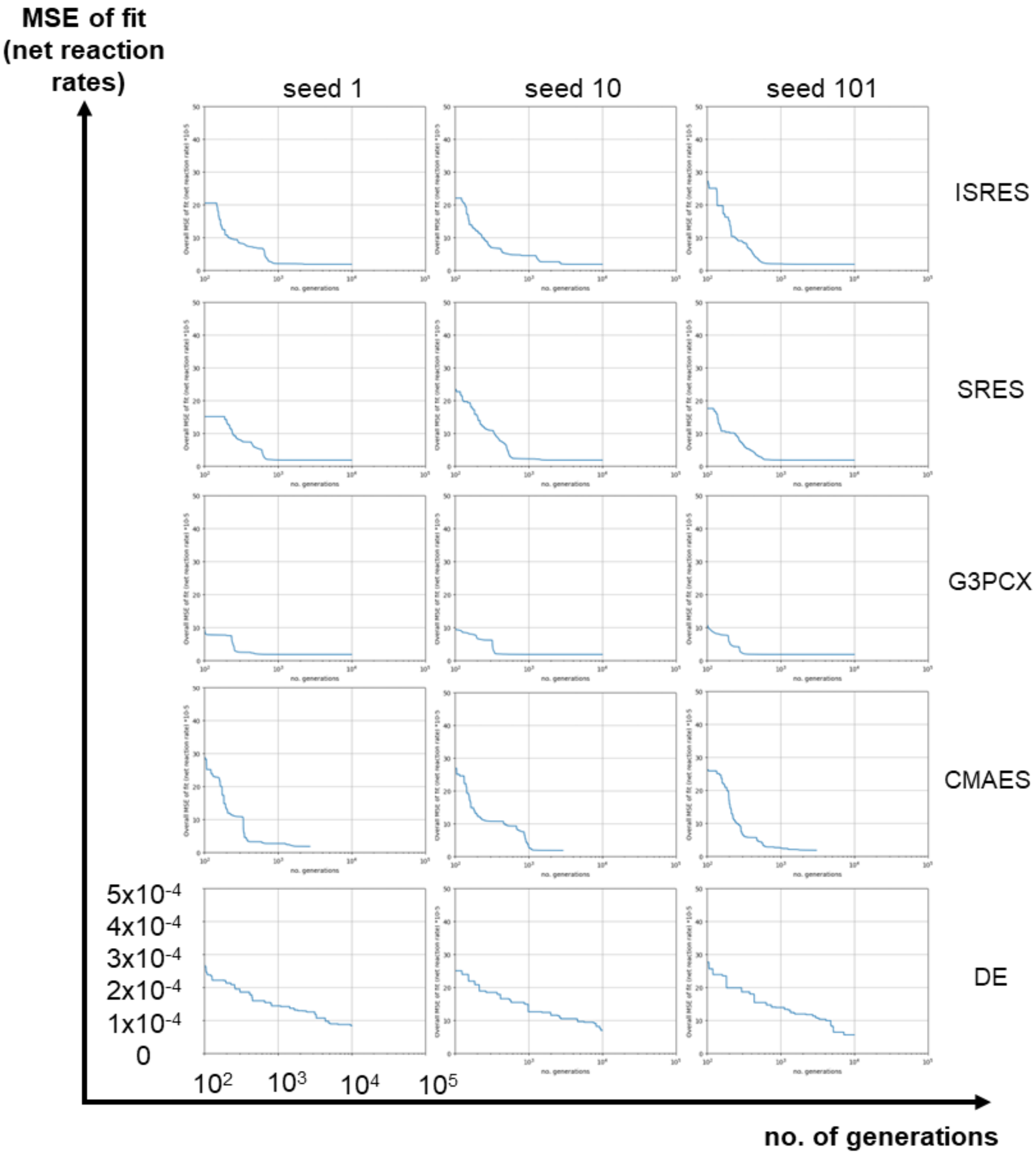


**Figure S2. Convergence profile in fitting the *Michaelis-Menten* model by candidate evolutionary algorithms and seed-instances.**

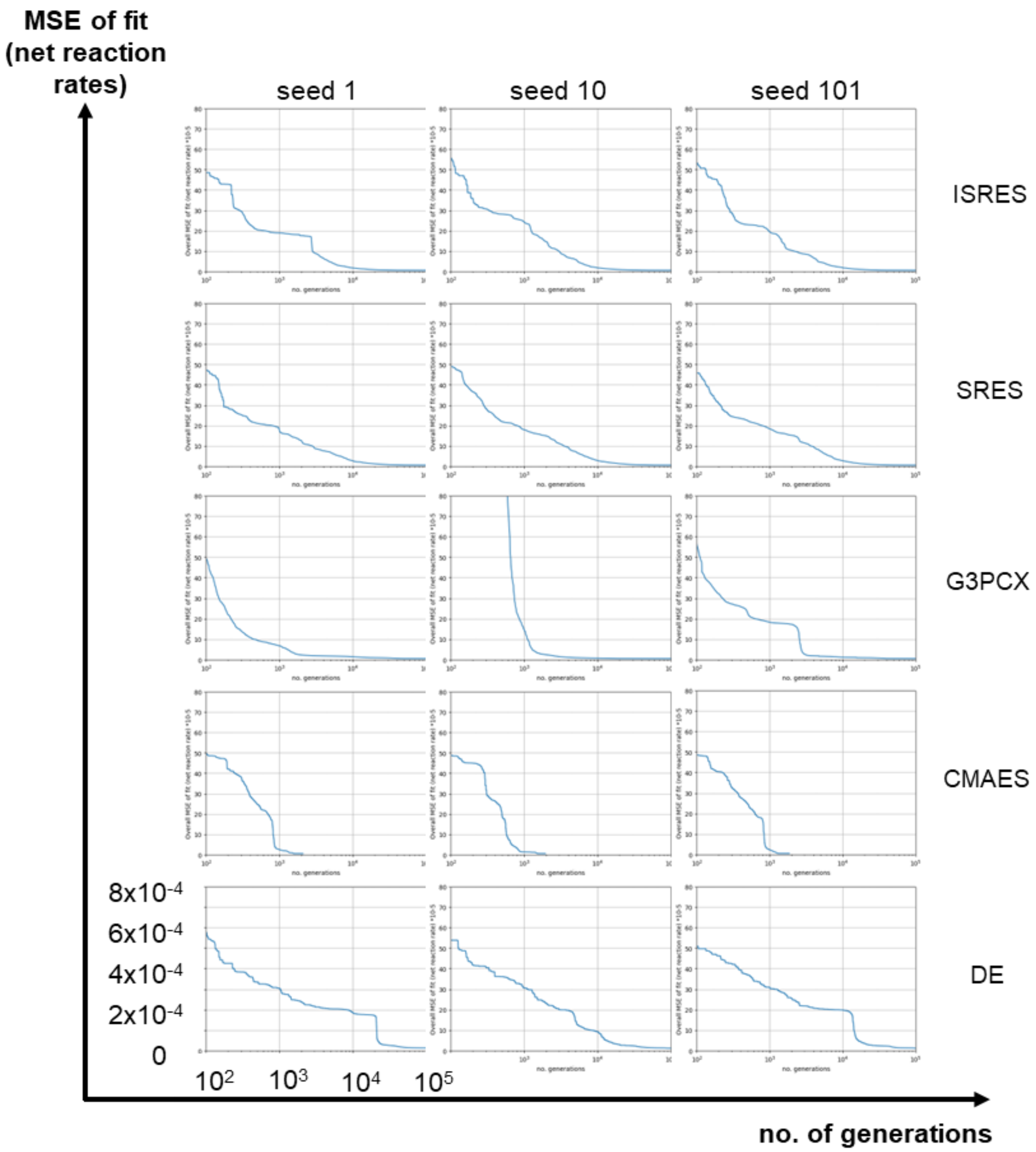


**Figure S3. Convergence profile in fitting the *Linlog* model by candidate evolutionary algorithms and seed-instances.**

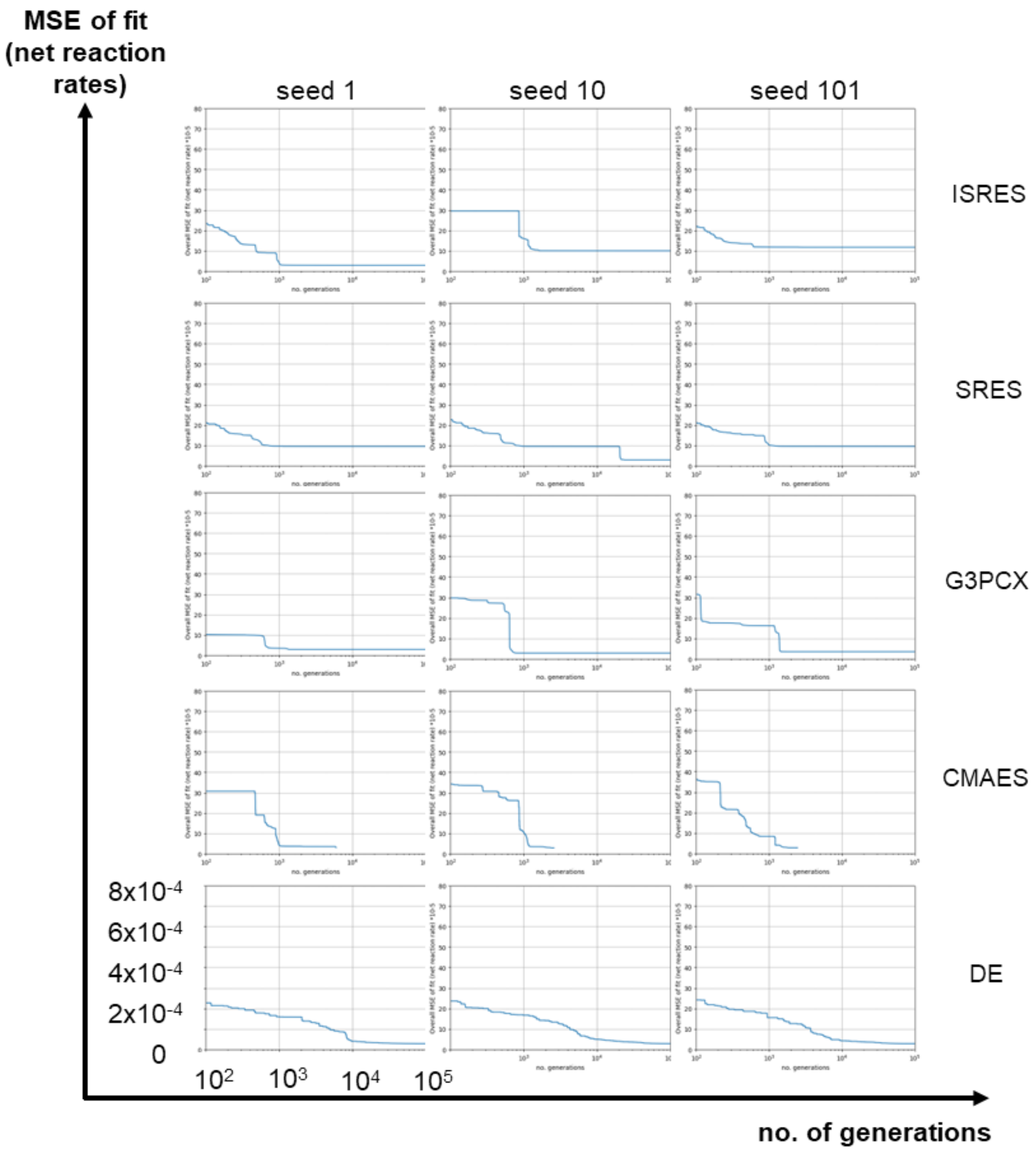


**Figure S4. Convergence profile in fitting the *convenience rate law* model by candidate evolutionary algorithms and seed instances.**

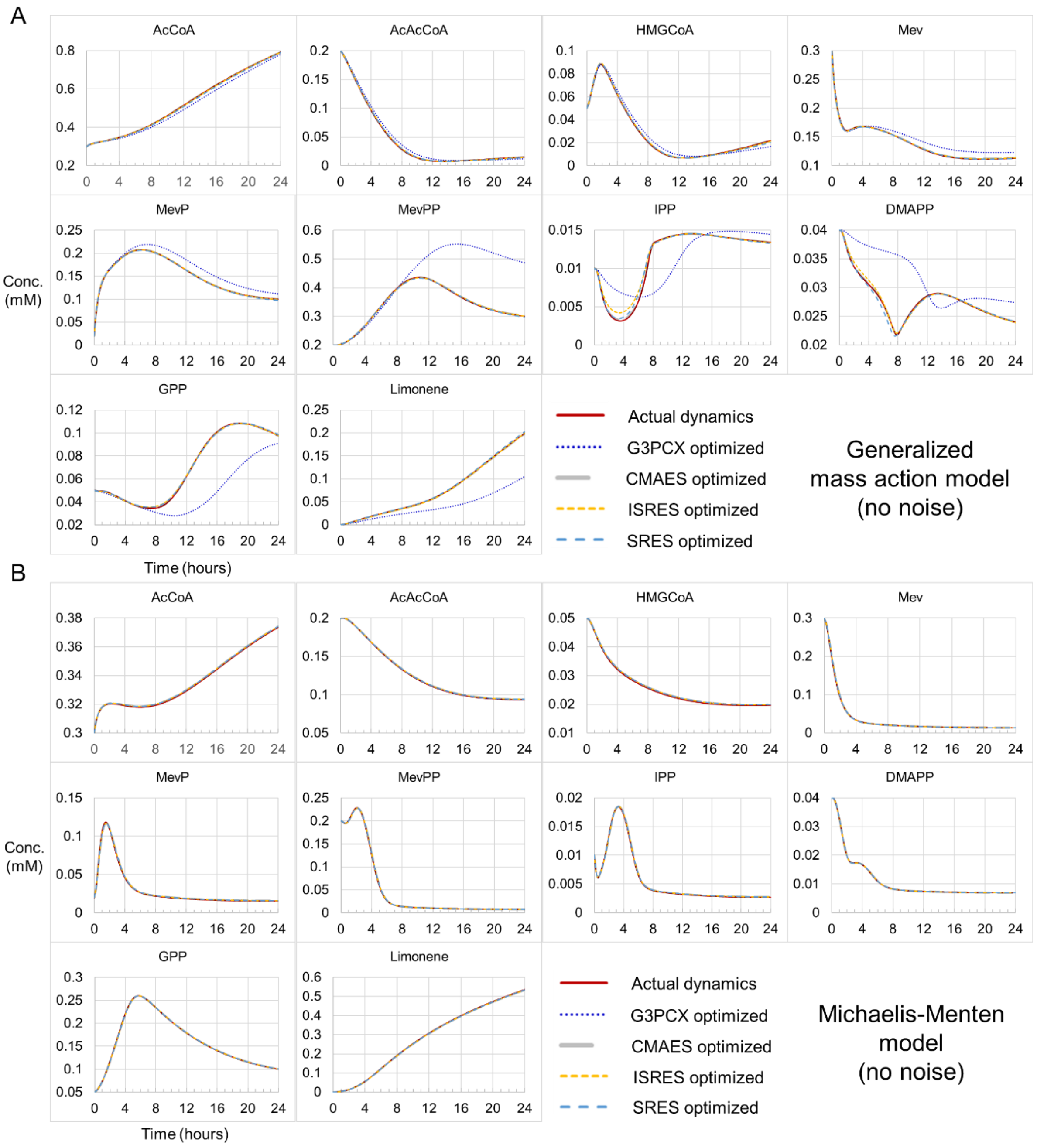


**Figure S5. Simulated generalized mass action (A) and Michaelis-Menten dynamics (B) based on parameters optimized using selected evolutionary algorithms for the case of zero measurement error.** The mean parameter values of 3 seed-instances are used.

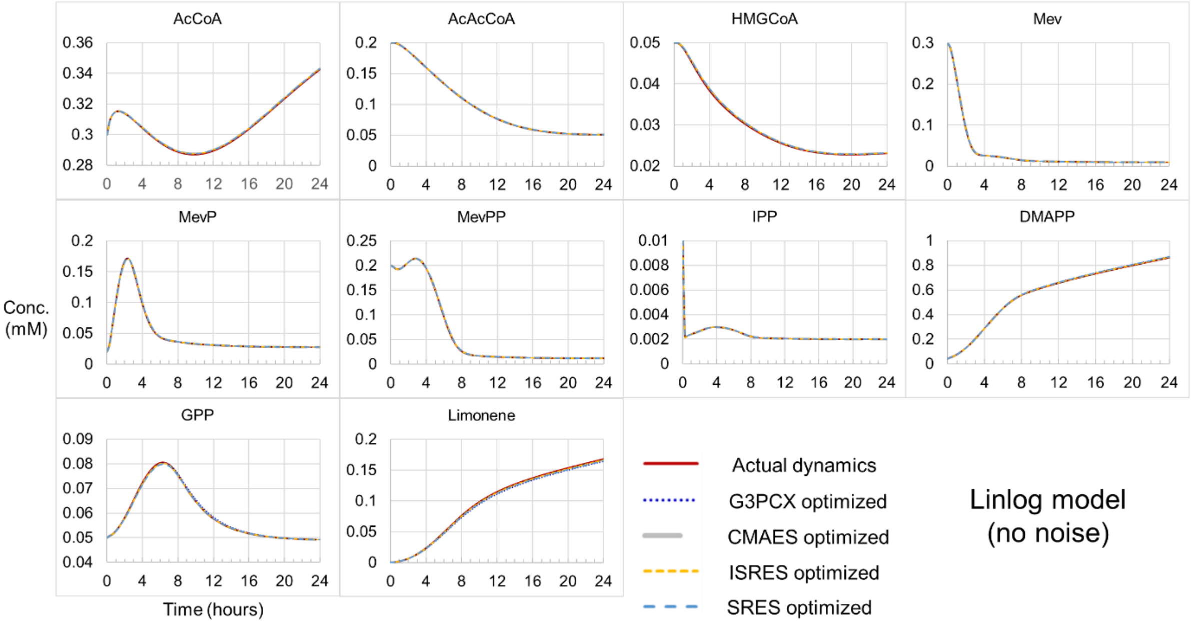


**Figure S6. Metabolite dynamics of Linlog model based on parameters optimized using selected evolutionary algorithms for the case of zero measurement error.** The mean parameter values of 3 seed-instances are used.

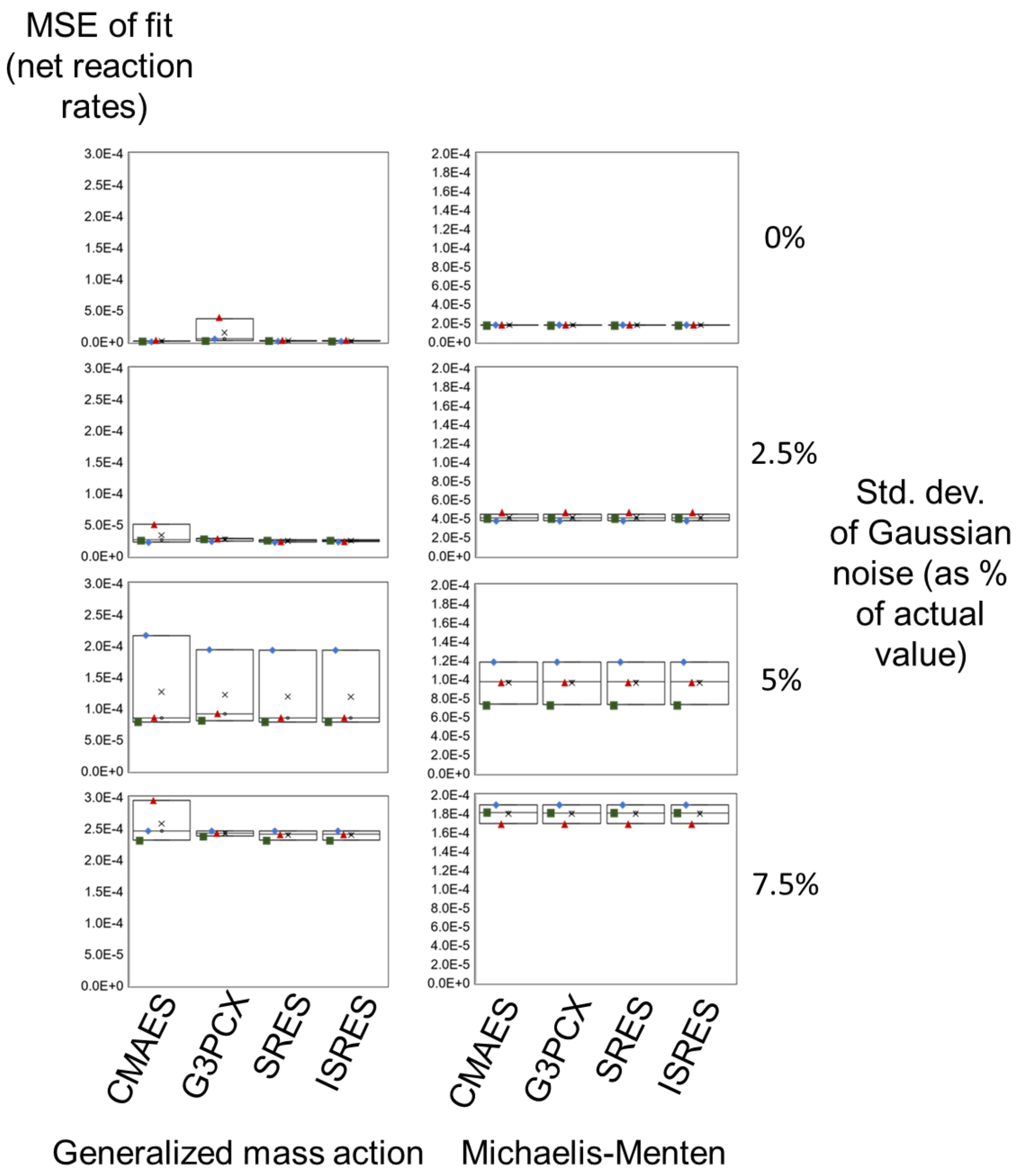


**Figure S7. Box plot distribution of the final cost function (MSE-of-fit) values for candidate evolutionary algorithms upon reaching the termination criterion.** Plots are shown for both generalized mass action and Michaelis-Menten models at various noise levels. Individual seed-instances and corresponding triplicate datasets are represented by squares, diamond shapes, and triangles. The horizontal lines of each box represent the 25%, 50%, & 75% of the distribution, while the cross indicates the mean value.

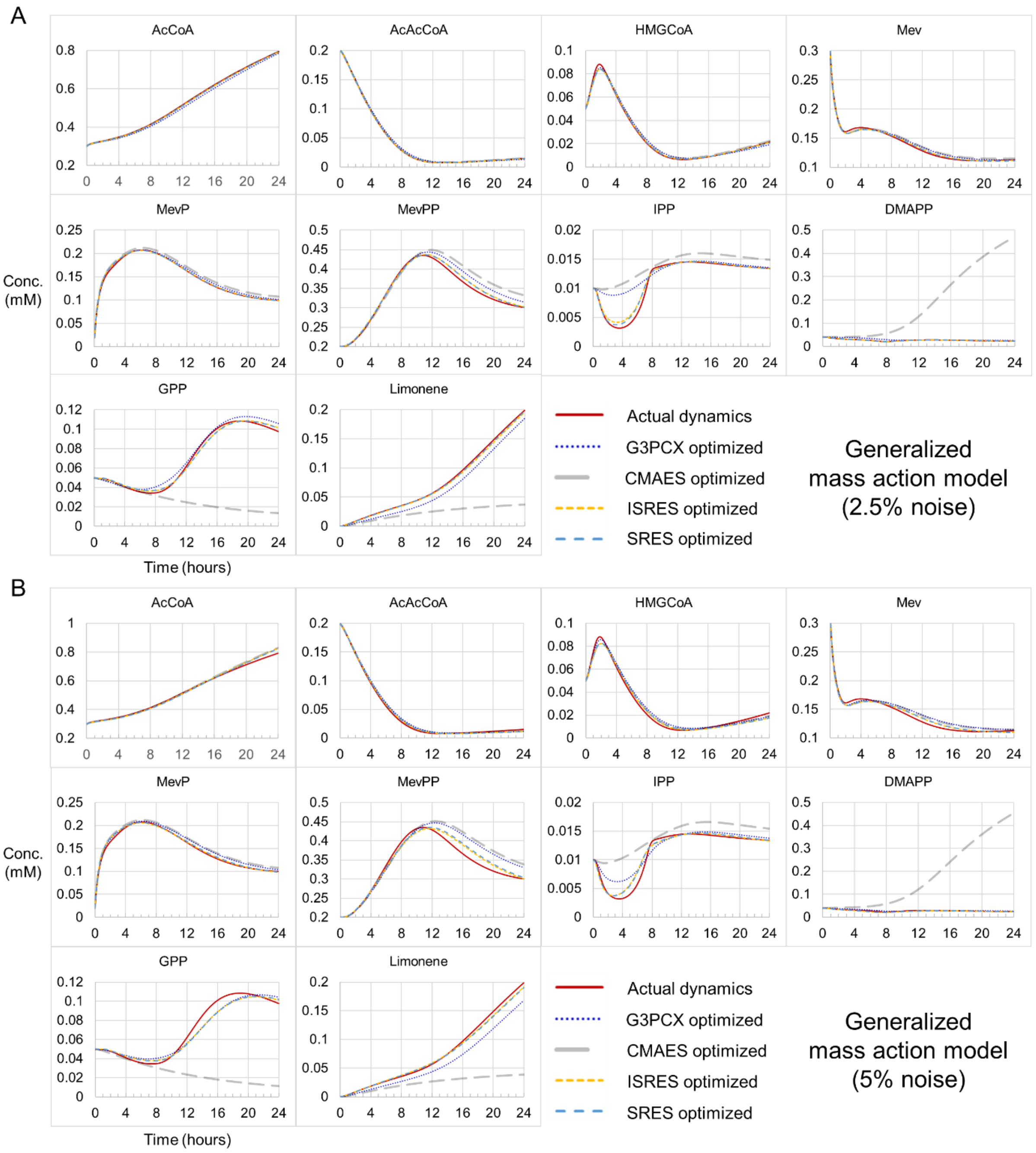


**Figure S8. Simulated mass action dynamics based on parameters estimated using selected evolutionary algorithms.** Results are shown for the case whereby measurement errors are sampled from a normal distribution centred on zero with a standard deviation equivalent to **A.** 2.5% and **B.** 5% of the underlying metabolite/enzyme concentration.

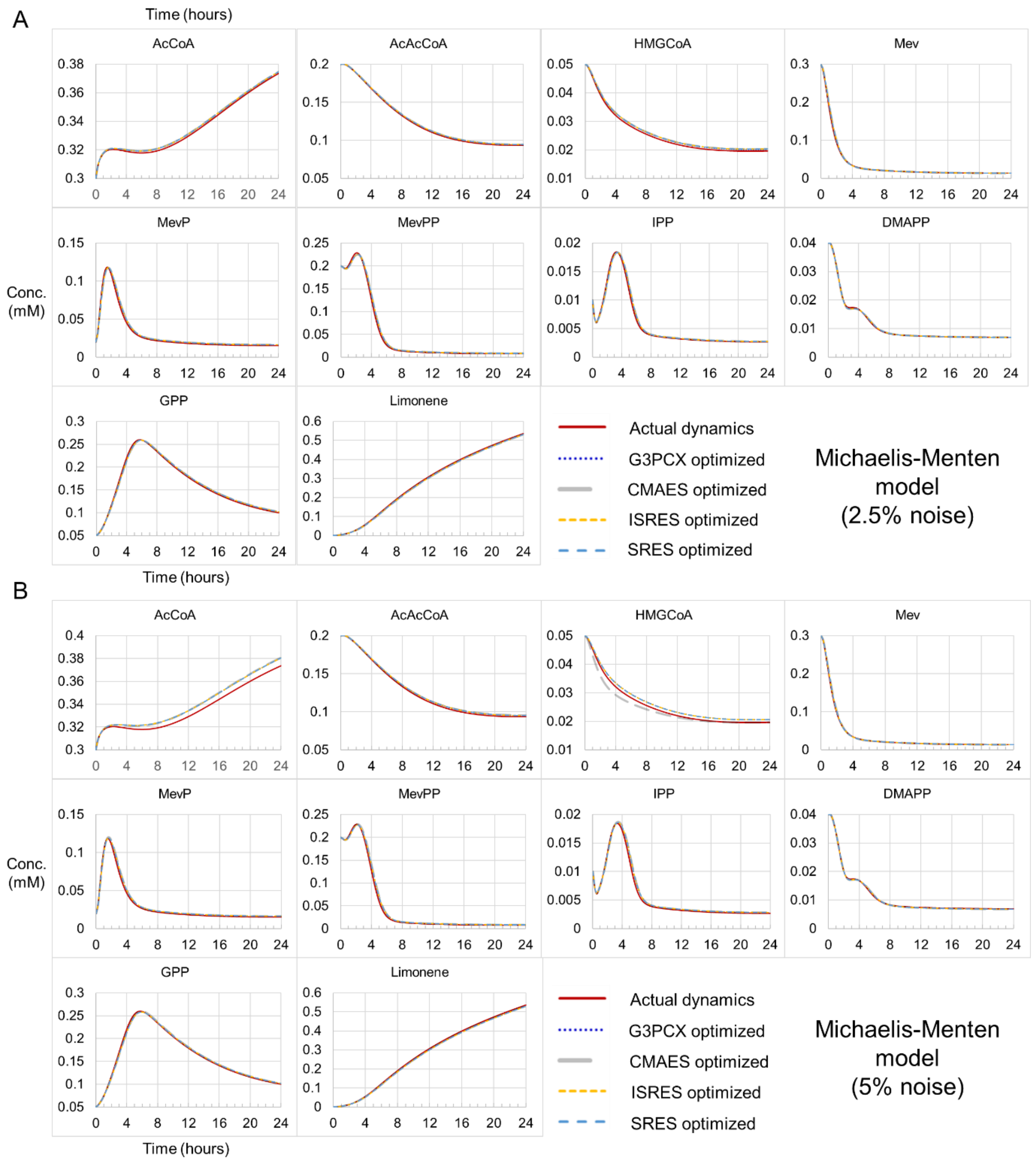


**Figure S9. Simulated Michaelis-Menten dynamics based on parameter values that are optimized using selected evolutionary algorithms.** Results are shown for the case whereby measurement errors are sampled from a normal distribution centred on zero with a standard deviation equivalent to **A.** 2.5% and **B.** 5% of the underlying metabolite/enzyme concentration.

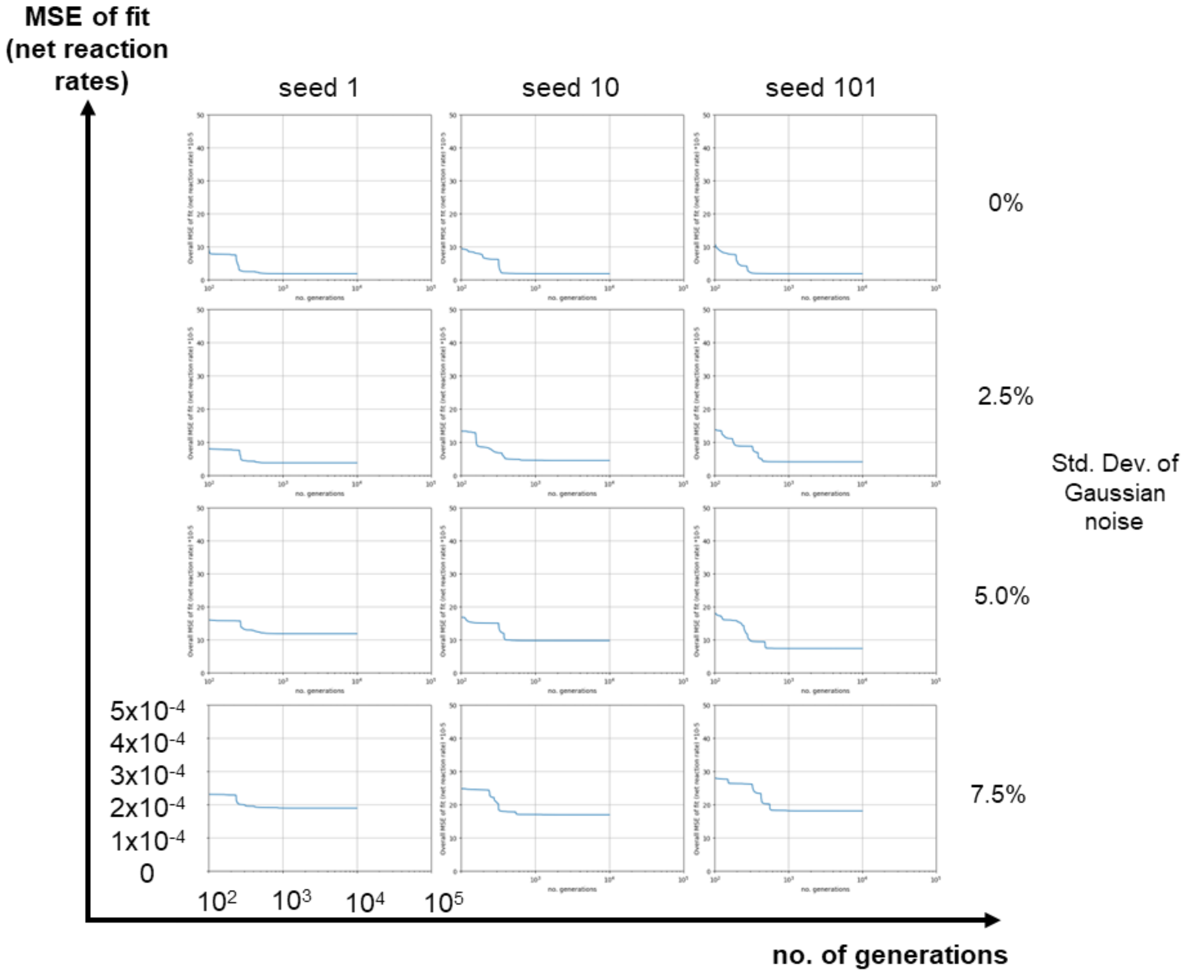


**Figure S10. Convergence profile in fitting the Michaelis-Menten model at various levels of Gaussian measurement noise using the G3PCX algorithm.**

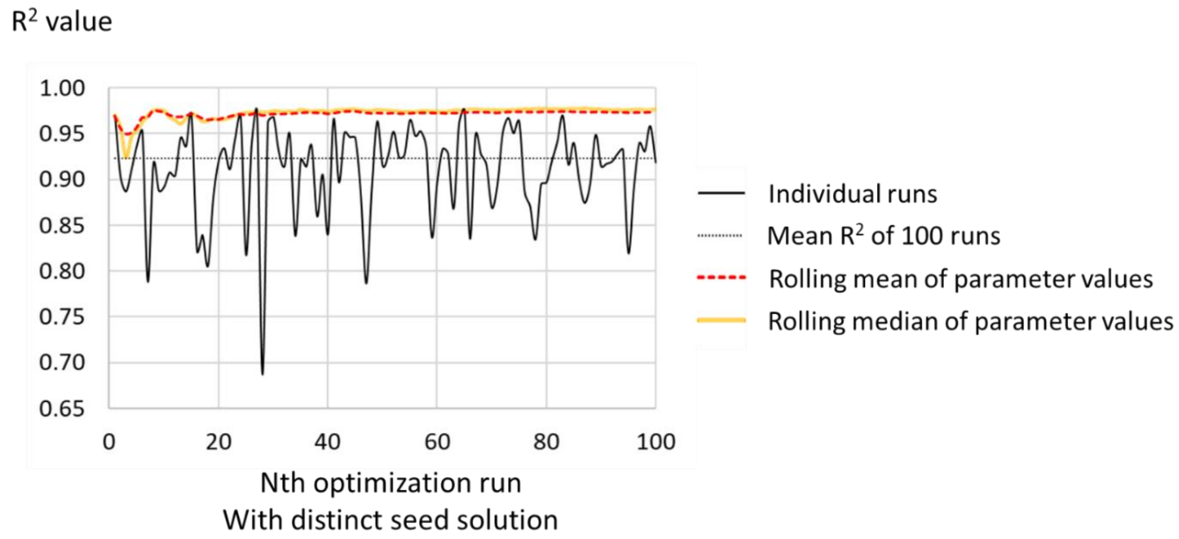


**Figure S11. Cumulative effect of individual optimizations on accuracy and reliability of parameter estimations.** As a proxy metric for accuracy, the resulting $R^2$ value from taking the mean/median of parameter estimations from individual seed solutions are compared cumulatively to that of the individual runs and their mean value. The result shown is for the case of G3PCX optimization of Michaelis-Menten kinetics at ‘5% noise level’.

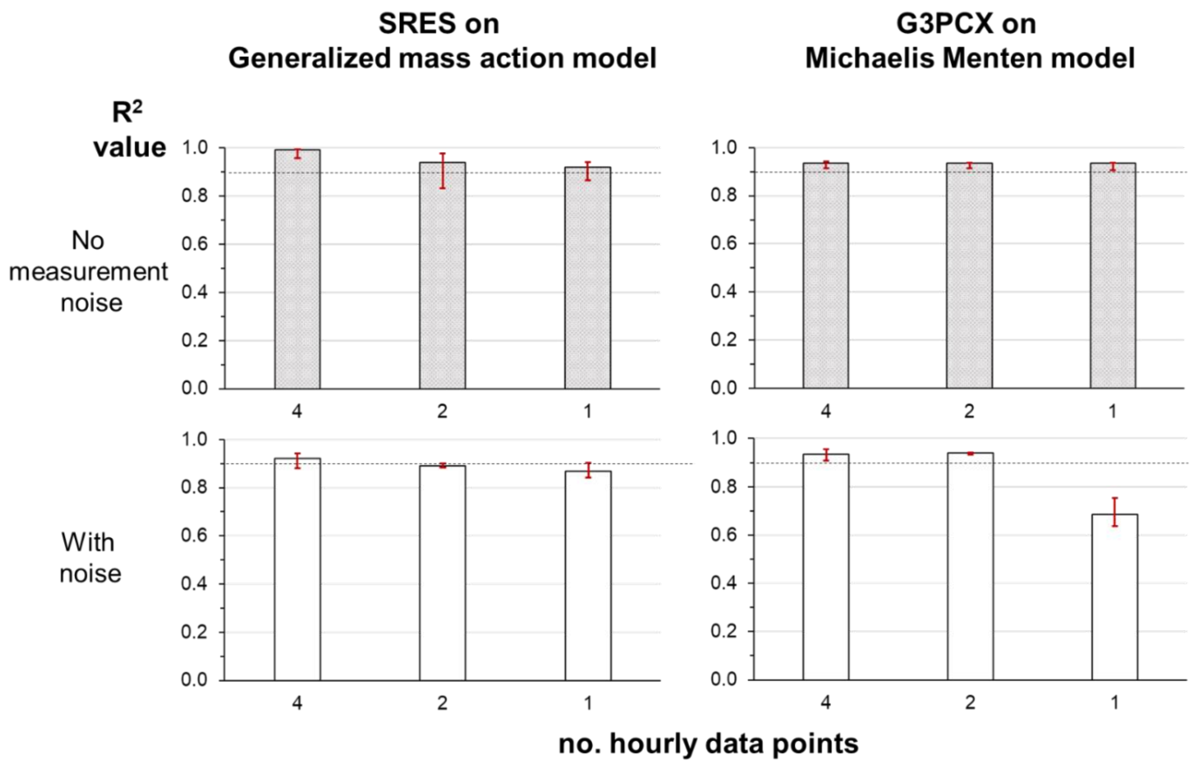


**Figure S12. The effect of the number of hourly data points on the quality of parameter predictions as reflected by the $R^2$ metric.** Two case studies are shown: estimation of generalized mass action (GMA) parameters using the SRES algorithm (left column) and the Michaelis-Menten (MM) parameter estimations utilizing the G3PCX algorithm (right column). The top row and bottom row are for the cases without and with measurement noise, respectively. For GMA parameter predictions, the amount of measurement noise is sampled from a normal distribution with mean zero and a standard deviation equivalent to 5% of the underlying metabolite/enzyme measurement. The corresponding value is 7.5% for MM parameter predictions. The highest noise level is chosen in each case such that the algorithm performs well with 4 hourly datapoints. The bars represent the median values while the lower and upper bounds (in red) mark the smallest and largest values based on three initial seed solutions.

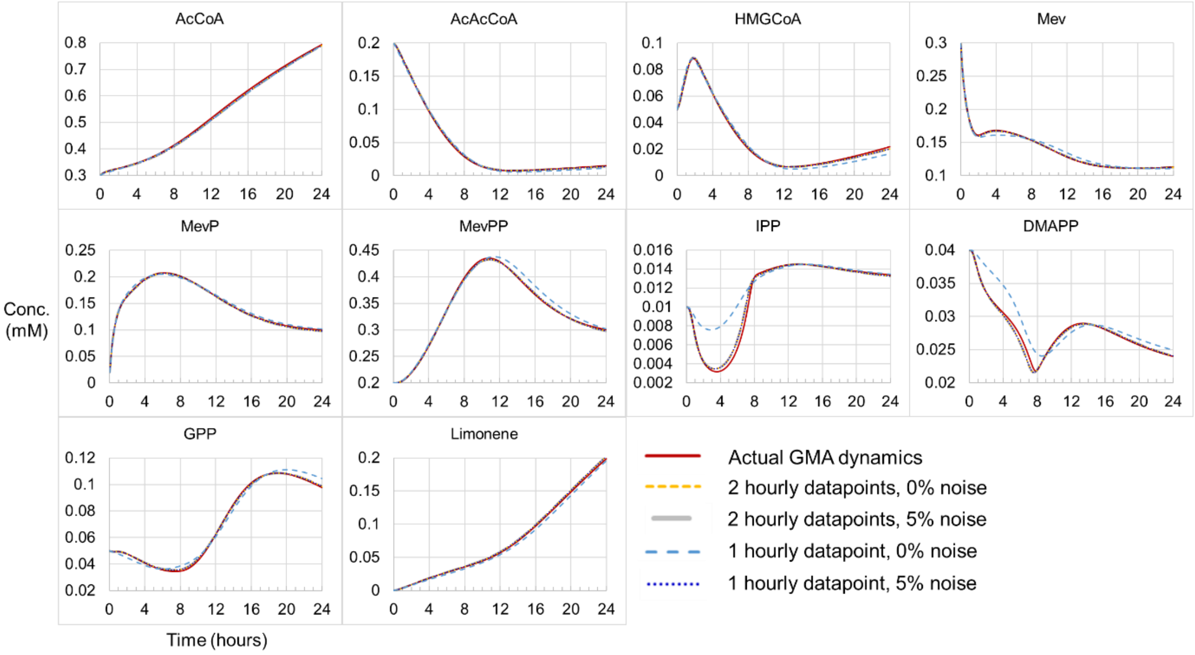


**Figure S13. Simulated generalized mass action dynamics based on parameter estimations using (metabolite and enzyme concentration) data with reduced hourly datapoints, i.e., two and one hourly datapoints.** The evolutionary algorithm in use is SRES, while four hourly datapoints were earlier used for the screening of evolutionary algorithms.

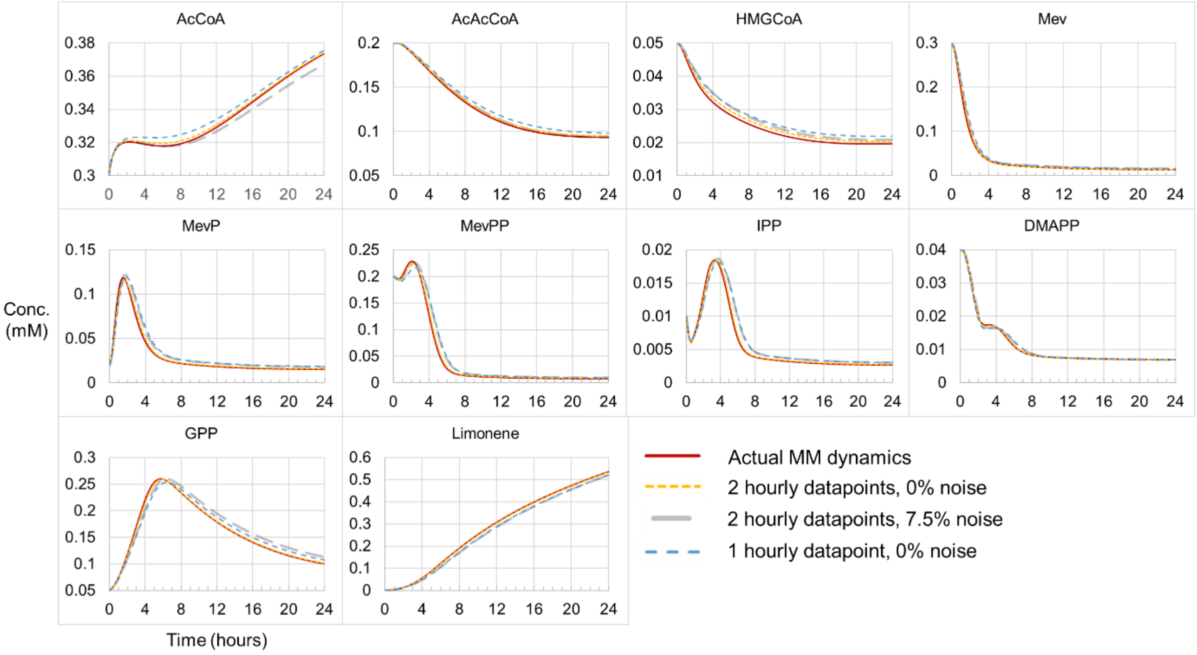


**Figure S14. Simulated Michaelis-Menten dynamics based on parameter estimations using (metabolite and enzyme concentration) data with reduced hourly datapoints, i.e., two and one hourly datapoints.** The evolutionary algorithm in use is G3PCX, while four hourly datapoints were earlier used for the screening of evolutionary algorithms.

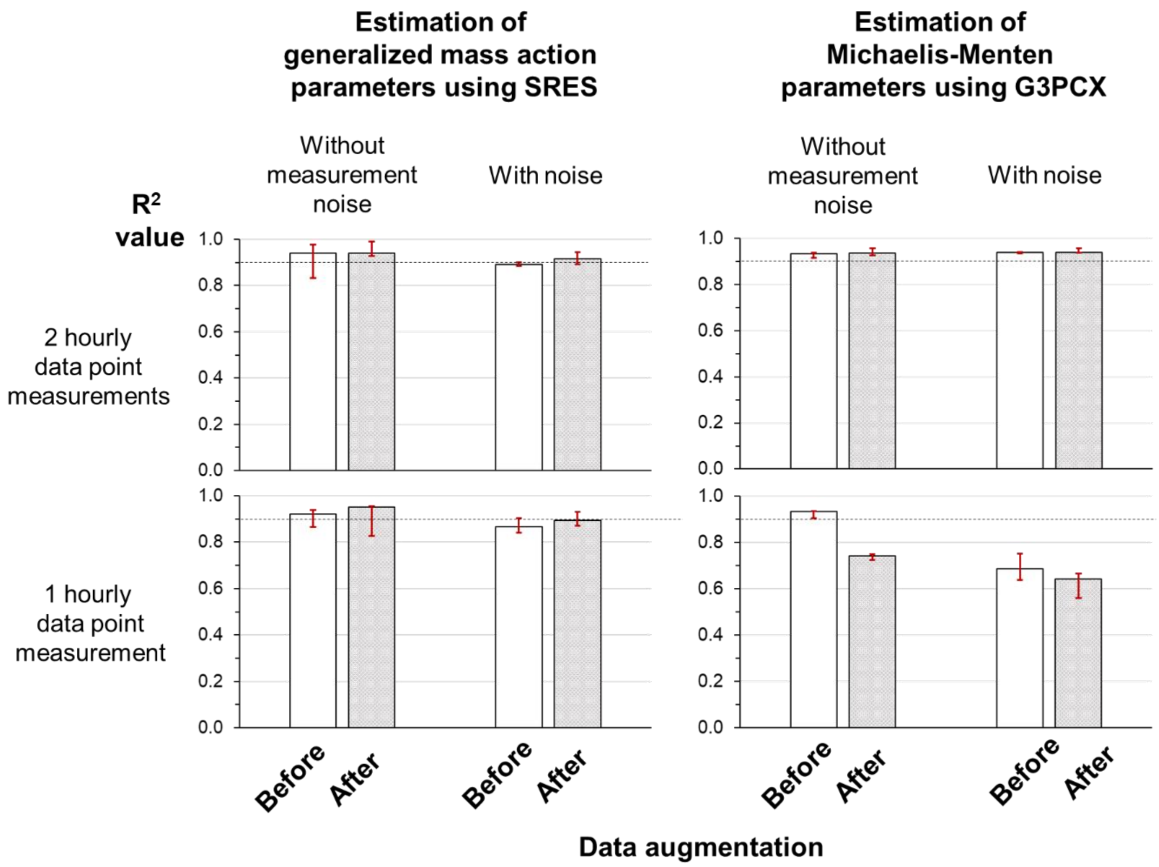


**Figure S15. The effect of data augmentation on the quality of parameter estimations as reflected by the $R^2$ metric.** Two case studies are shown: estimation of generalized mass action (GMA) parameters using the SRES algorithm (left column) and the Michaelis-Menten (MM) parameter estimations utilizing the G3PCX algorithm (right column). Data augmentations (Materials & methods) are done in two cases to increase the hourly data points from 2 (top row) and 1 (bottom row) to 4 per hour. For GMA parameter predictions, the amount of measurement noise is sampled from a normal distribution with mean zero and a standard deviation equivalent to 5% of the underlying metabolite/enzyme measurement. The corresponding value is 7.5% for MM parameter predictions. The bars represent the median values while the lower and upper bounds (in red) mark the smallest and largest values based on three initial seed solutions.

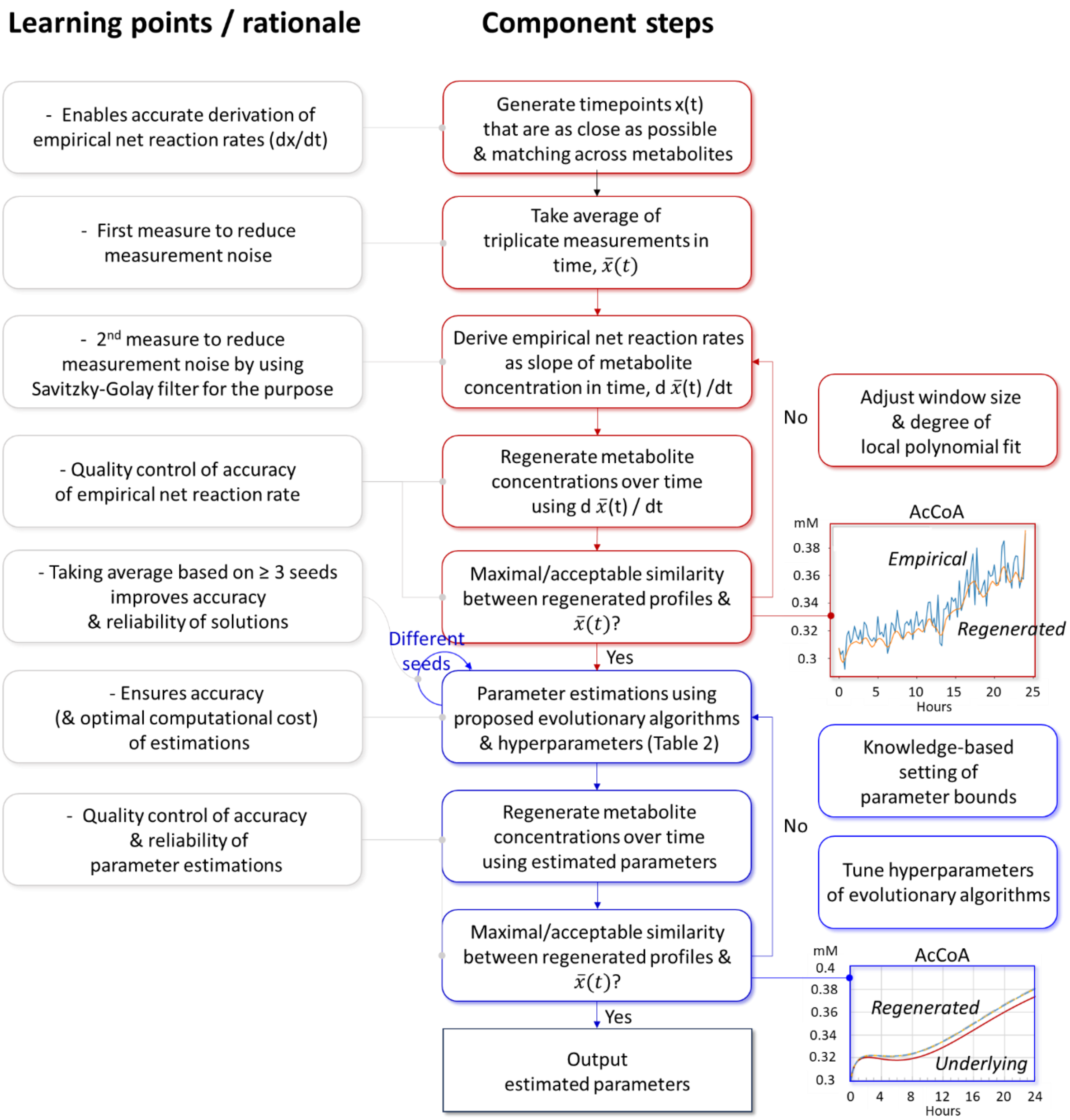


**Figure S16. Learning points and approach for effective estimation of kinetic parameters.** The grey boxes outline the rationale or learning point of associated steps. The red boxes depict the 5 preprocessing steps to derive high quality values for empirical net reaction rates, while the blue boxes set out the three subsequent steps for parameter estimations.
